# Engineering amino acid-derived malonyl-CoA pathways to boost polyketide production in *Yarrowia lipolytica*

**DOI:** 10.1101/2025.08.03.668335

**Authors:** Jinpeng Wang, Yuxiang Hong, Zizhao Wu, Ayelet Fishman, Peng Xu

## Abstract

Malonyl-CoA is a central precursor involved in the synthesis of various bio-based chemicals, including polyketides, fatty acids, and flavonoids. However, the production of these chemicals is often limited by the availability of malonyl-CoA. Based on retrosynthesis principles, we designed two thermodynamically favorable malonyl-CoA pathways using L-glutamate and L-aspartate as substrates. The novel pathways leverage oxidative deamination and decarboxylation reactions and efficiently channel metabolic flux toward malonyl-CoA, resulting in increased production of total polyketides beyond the capacity of the native acetyl-CoA carboxylase route using glucose as substrate. We also discovered a new-to-nature polyketide (4-hydroxy-6-hydroxyethyl-2-pyrone) derived from the side activity of the TAL pathway, reaching 6.4 g/L in *Y. lipolytica*. This work highlights the utility of the novel malonyl-CoA pathways in enhancing polyketide production, and the possibility of upcycling abundant amino acids or protein waste in the animal farming or meat industry to produce high-value nonnatural polyketides.

**Highlights:**

- Two thermodynamically favorable malonyl-CoA pathways in *Yarrowia lipolytica* using L-glutamate and L-aspartate as substrates were engineered.
- The novel oxygen-dependent malonyl-CoA pathways outperformed the canonical acetyl-CoA carboxylase pathway in polyketide production.
- A new-to-nature polyketide, 4-hydroxy-6-hydroxyethyl-2-pyrone (HHEP), was produced at a remarkably high titer of 6.4 g/L in shaking flask.
- The engineered pathway improved the total production of malonyl-CoA-derived compounds (TAL, HHEP, and lipids).

## 1 Introduction

Malonyl-CoA is the C_2_ building block in the biosynthetic pathway of fats (*1, 2*), oleochemicals (*3*), and polyketides (*4-6*) in nature. As such, it sits at the heart of central metabolism in both prokaryotes and eukaryotes and underpins a vast range of biochemical transformations. Beyond its well-known role in fatty acid biosynthesis, malonyl-CoA is also integral to the biosynthesis of numerous high-value compounds including flavonoids (*7, 8*) and many pharmaceutically relevant polyketides (*9-13*). Consequently, boosting intracellular malonyl-CoA availability has become a central goal in metabolic engineering (*14, 15*).

Given malonyl-CoA’s central importance, the quest for efficient and feasible malonyl-CoA pathways has long been a focus of the metabolic engineering community. The canonical route relies on pyruvate dehydrogenase (PDH) converting pyruvate into acetyl-CoA, followed by acetyl-CoA carboxylase (ACC) converting acetyl-CoA into malonyl-CoA (**Fig. 1**) (*16*). Although widely employed in industrial microorganisms, this PDH–ACC pathway carries a moderate overall negative Gibbs free energy (Δ_r_G^m^ ⍰ ≈ –51 kJ/mol) yet also entails high ATP consumption (**Table 1**). A second strategy uses malonyl-CoA synthetase (MCS) (*17-19*) to directly activate malonate into malonyl-CoA (Δ_r_G^m^ ⍰ ≈ –53.4 kJ/mol), avoiding the carboxylation step and offering kinetic advantages due to simpler enzyme regulation. However, the cost of malonate is economically prohibitive (8,000$/ton). This route also imposes the cells high consumption of ATP. Recently, an alternative “transaminase–malonyl-CoA reductase” route (Fig. 1) was introduced to improve polyketide synthesis (*20*) in *E. coli*, wherein malonate semialdehyde (arising via transamination from β-alanine) is converted to malonyl-CoA by a specialized malonyl-CoA reductase. Although Gibbs energy for this emerging pathway (Δ_r_G°^m^⍰ ≈ –16 kJ/mol for the enzyme cascade) can be favorable under standard conditions (species concentration of 1 M), the standard Gibbs free energy under physiologically-relevant conditions (Δ_r_G^m^ ⍰ ≈ +0.6 kJ/mol, species concentration of 1 mM) is thermodynamically unfavorable. In addition, existing enzyme pairs of transaminase and reductase demonstrated mediocre *K*_m_ and *k*_cat_ values. This necessitates us to find alternative malonyl-CoA pathways to improve microbial production of important metabolites.

**Table 1.** Summary of the reaction components of canonical (**a, b**), noncanonical (**c**) and the new predicted malonyl-CoA (**d** and **e**) pathways.

| Pathway | Reaction | Cofactor | ATP<br>consume | CO<sub>2</sub><br>release | Redox<br>balance | Enzyme | EC | Source |
| --- | --- | --- | --- | --- | --- | --- | --- | --- |
| Canonical | a<br>Pyruvate + CoASH + NAD <=> Acetyl-CoA + NADH + CO<sub>2</sub> | CoASH,NAD+ | 0 | 1 | NADH | PDH | 1.2.1.51 | Tong, L.<br>(2005) ( 16) |
|  | Acetyl-CoA + ATP + CO<sub>2</sub> + H<sub>2</sub>O <=> ADP + Malonyl-CoA + Pi | NA | 1 | 0 | 0 | ACC | 6.4.1.2 |  |
|  | b<br>Malonate + CoASH + ATP +H<sub>2</sub>O <=> AMP + Malonyl-CoA + 2Pi | CoASH | 2 | 0 | 0 | MS | 6.2.1.76 | Wakil, S. J.<br>(1958) ( 19) |
| Noncanonical | c<br>beta-Alanine + Pyruvate <=> alpha-Alanine + Malonate semialdehyde | NA | 0 | 0 | 0 | TA(BauA) | 2.6.1.18 | Li, J., et al.<br>(2024) (20) |
|  | Malonate semialdehyde + CoASH + NADP <=> Malonyl-CoA + NADPH | CoASH,NADP+ | 0 | 0 | NADPH | MCR-C | 1.2.1.75 |  |
| Predicted | d<br>L-glutamate + 2-oxoglutarate + O<sub>2</sub> <=> 3-hydroxy-L-glutamate + succinate + CO<sub>2</sub> | NA | 0 | 1 | 0 | IboH | 1.14.11.76 | Obermaier,<br>S., et al.<br>(2020)(21),<br>Nie, M., et<br>al. (2024)<br>(22) |
|  | 3-Hydroxy-L-glutamate <=> Malonate semialdehyde + Glycine | NA | 0 | 0 | 0 | LtaE | 4.1.2.48 |  |
|  | Malonate semialdehyde + CoASH + NADP <=> Malonyl-CoA + NADPH | CoASH,NADP+ | 0 | 0 | NADPH | MCR-C | 1.2.1.75 |  |
|  | e<br>Aspartate <=> beta-Alanine + CO<sub>2</sub> | NA | 0 | 1 | 0 | ADC | 4.1.1.11 | Cronan, J.,<br>et al.<br>(1980)(23),<br>Yamashita,<br>M., et al.<br>(1996) (24) |
|  | beta-Alanine + H<sub>2</sub>O+ O<sub>2</sub> <=> Malonate semialdehyde + H<sub>2</sub>O<sub>2</sub> + NH<sub>3</sub> | NA | 0 | 0 | 0 | MAO | 1.4.3.21 |  |
|  | Malonate semialdehyde + CoASH + NADP <=> Malonyl-CoA + NADPH | CoASH,NADP+ | 0 | 0 | NADPH | MCR-C | 1.2.1.75 |  |

**Fig. 1.**
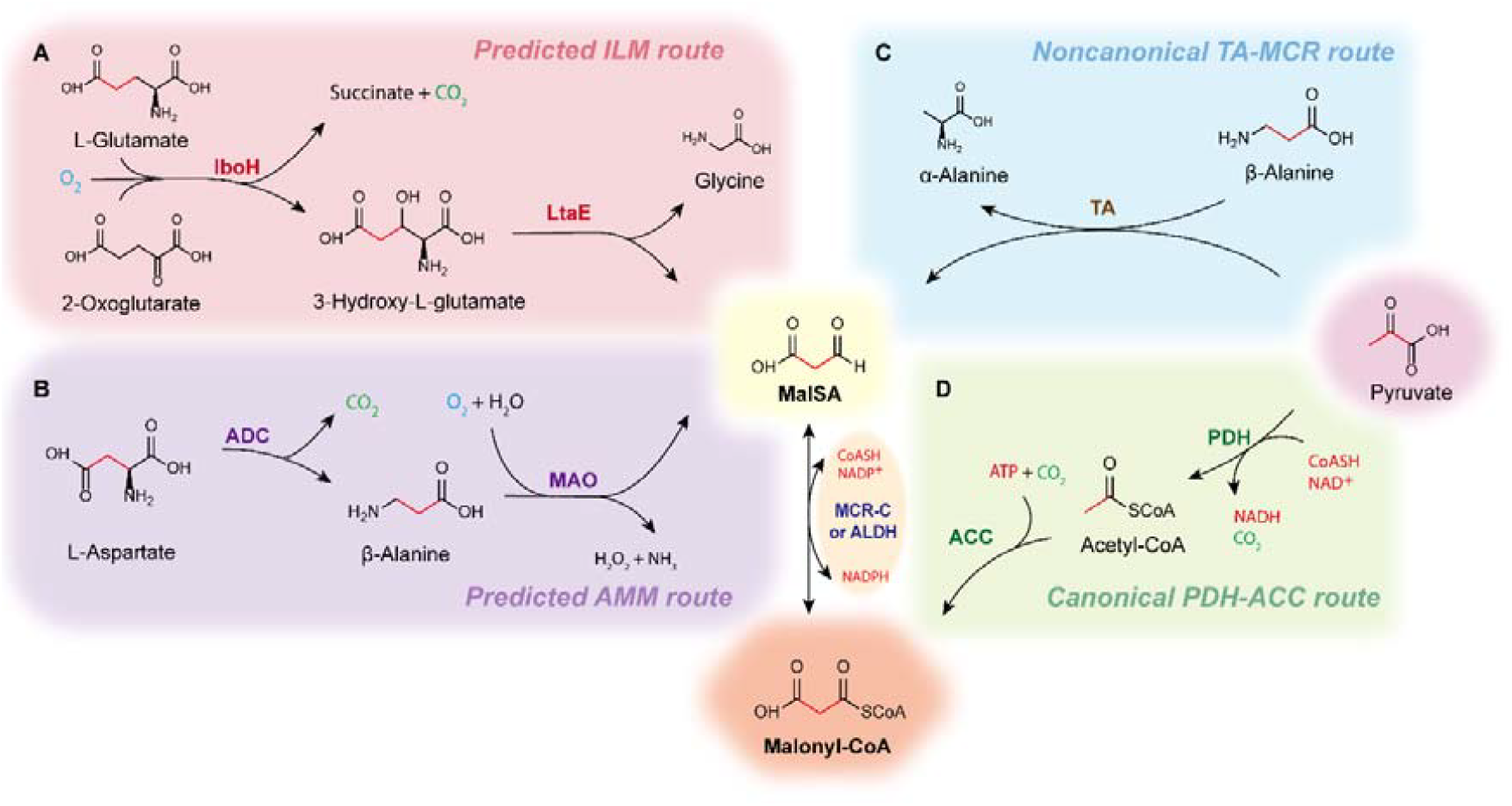
Malonyl-CoA canonical, noncanonical, and predicted pathways: (**A**) Novel O_2_-dependent malonyl-CoA pathways derived from L-glutamate (ILM route). IboH: 2-oxoglutarate-dependent L-glutamate dioxygenase; LtaE: aldolase; MCR-C: malonyl-CoA reductase C-terminal; (**B**) Novel O_2_-dependent malonyl-CoA pathways derived from L-aspartate (AMM route). ADC: aspartate decarboxylase, MAO: monoamine oxidase; (**C**) Noncanonical TA-MCR route involves malonate semialdehyde (MalSA). TA: transaminase; (**D**) Canonical PDH-ACC route. PDH: pyruvate dehydrogenase, ACC: acetyl-CoA carboxylase.

An ideal malonyl-CoA pathway should offer greater thermodynamic favorability, improved catalytic constants, or less stringent cofactor and ATP requirements. In this study, we addressed this challenge by retro-synthetically designing and implementing two novel malonyl-CoA routes, starting from the abundant amino acids L-glutamate and L-aspartate (Fig. 1). Both pathways leverage malonate semialdehyde as the intermediate and malonyl-CoA reductase as a key step. Using the simplest polyketide TAL as a test molecule, we experimentally validated that these new pathways efficiently redirect carbon flux toward malonyl-CoA and boosted TAL and a new-to-nature (4-hydroxy-6-hydroxyethyl-2-pyrone) polyketide production in *Y. lipolytica*. Our re-designed malonyl-CoA pathways expanded the biotechnological repertoire of high-value polyketides from low-cost and abundant amino acids.

## 2. Results and discussion

### 2.1 Designing novel oxygen-dependent malonyl-CoA pathways from two abundant amino acids

We began by applying a retrosynthetic framework to identify possible malonyl-CoA pathways from two abundant amino acids, L-glutamate and L-aspartate. Both proposed pathways generate malonate semialdehyde (MalSA) as an intermediate and then employ malonyl-CoA reductase (MCR) to convert MalSA to malonyl-CoA (**Fig. 1A** and **1B**). In the L-glutamate route, L-glutamate dioxygenase (IboH) (*21*) hydroxylates L-glutamate to form 3-hydroxy-L-glutamate, which subsequently undergoes an aldol cleavage (*22*), yielding MalSA and glycine. In the L-aspartate route, aspartate decarboxylase (ADC) converts L-aspartate to β-alanine (*23*), then β-alanine undergoes an oxidative deamination to form MalSA by a monoamine oxidase (MAO) (*24*), generating hydrogen peroxide (H_2_O_2_) as byproduct. Both enzymes rely on oxidative deamination and decarboxylation steps releasing CO_2_, amine (or glycine) or H_2_O_2_, proceeding with increased entropy or disorder, signifying a strong driving force or large negative Gibbs energy change.

With an AI-powered kinetic data prediction tool UniKP (*25*) including reported kinetic data, and biochemical reaction thermodynamic calculator *eQuilibrator 3*.*0* (*26*), we retrieved the kinetic and thermodynamic information of the novel malonyl-CoA pathways. Compared to the canonical PDH-ACC and MCS routes, both the L-glutamate and L-aspartate pathways exhibit reduced ATP demand (**Table 1**), thereby imposing less cellular energy cost. Thermodynamic and kinetic analyses (**Table 2**) indicate that although both pathways rely on MCR — known to exhibit marginal thermodynamic favorability on its own, the large driving force by coupling with oxidative deamination and decarboxylation steps will thermodynamically benefit MCR. Specifically, IboH-aldolase (Δ_r_G ^m^⍰ ≈ -466.6 kJ/mol) or ADC-MAO (Δ_r_G^m^ ⍰≈ -146.3 kJ/mol) reactions each display large negative Gibbs free energy changes (**Table 2**), which collectively drive the overall pathway into a highly exergonic regime even when the MCR step by itself is relatively less favorable (*27-29*). This thermodynamic synergy sets our proposed pathways apart from the previously reported transaminase-MCR approach (*20*), whose net Gibbs energy under physiological conditions becomes slightly positive (Δ_r_G^m^ ⍰ ≈ +0.6 kJ/mol). Moreover, predicted *K*_m_ and *k*_cat_ values for IboH, aldolase, MAO, and MCR, fall within ranges better suitable for metabolic engineering (**Table 2**), than the transaminase-MCR route (*20*).

**Table 2.**
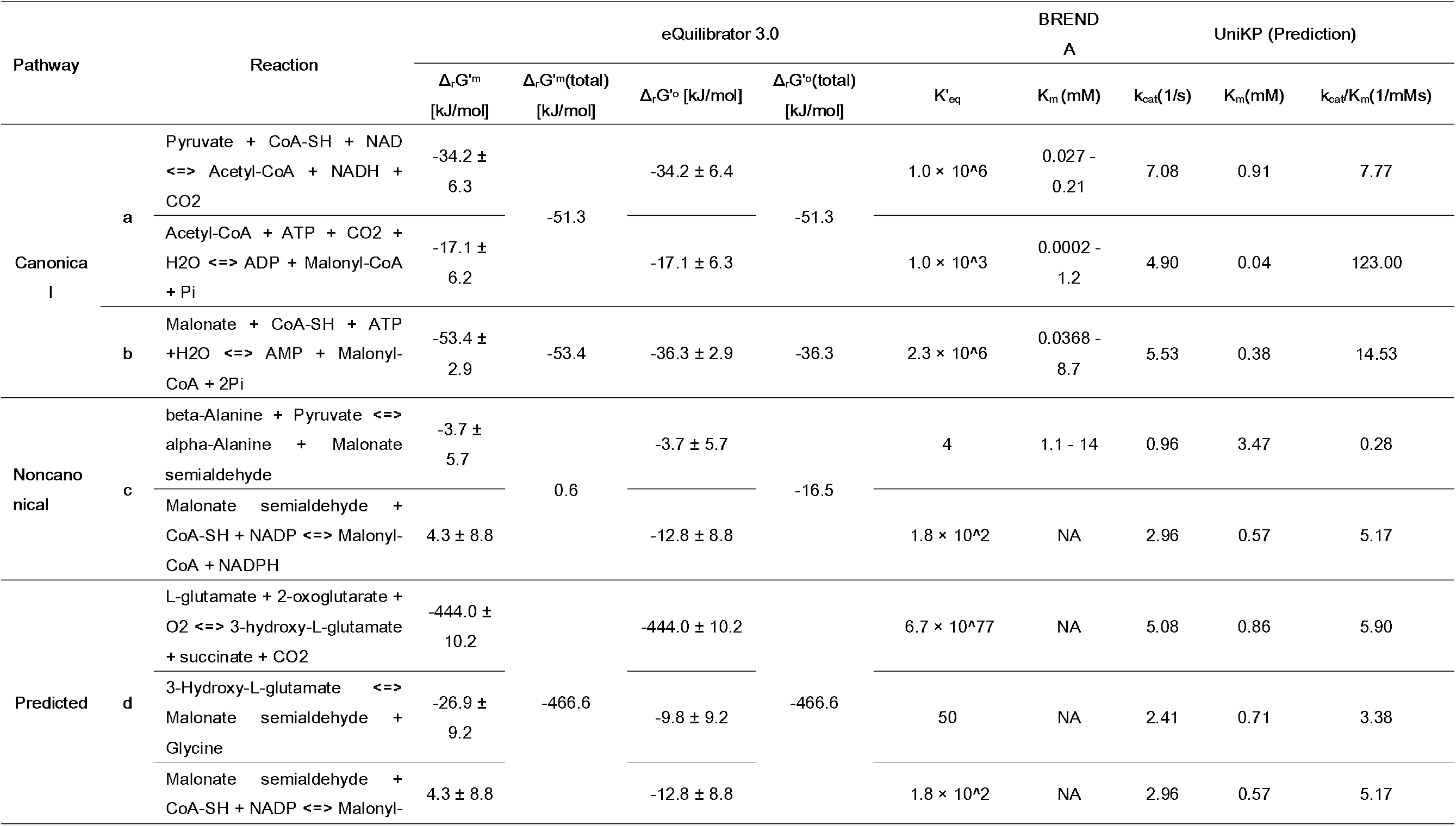
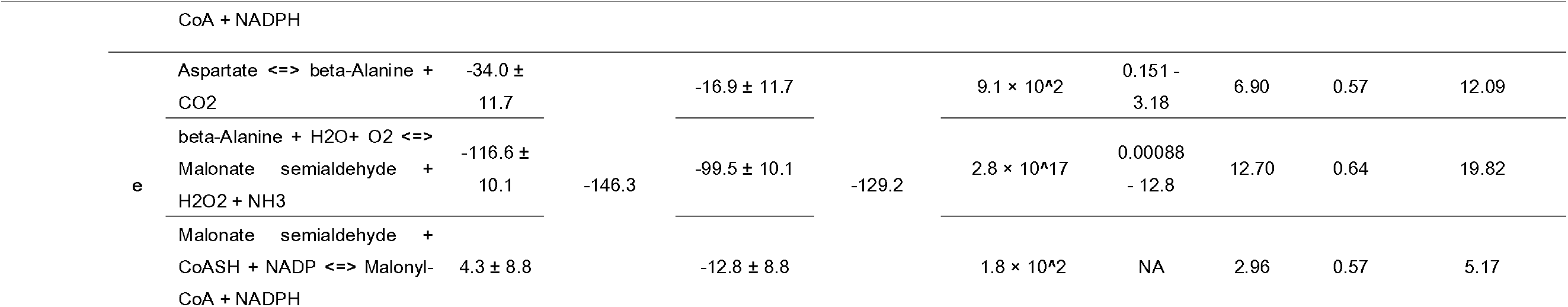
Summary of thermodynamic and kinetic parameters of canonical, noncanonical and the novel malonyl-CoA pathways.

### 2.2 Novel malonyl-CoA pathways boost TAL production and produce an unknown metabolite

We next validated whether the novel malonyl-CoA pathways could enhance malonyl-CoA level. Since intracellular malonyl-CoA or its intermediate MalSA are highly unstable, prone to rapid fluctuations, and tightly regulated within central carbon metabolism, we instead employed more stable malonyl-CoA derivatives to assess the activity of these two pathways. Here, Y. lipolytica was chosen as the host strain due to its naturally high malonyl-CoA flux, which favors the biosynthesis of polyketides and lipids. Therefore, the simplest polyketide (triacetic acid lactone) was chosen as the test molecule. We integrated a codon-optimized 2-pyrone synthase (*Gh*2PS) from *Gerbera hybrida* into the *lip1* locus of *Y. lipolytica* Po1fk (*30, 31*). This enzyme catalyzes the condensation of malonyl-CoA with acetyl-CoA (**Fig. 2A**) to form triacetic acid lactone (TAL), serving as a direct reporter for the newly designed malonyl-CoA pathways. The Lip1 gene, encoding a lipase involved in triglyceride hydrolysis, was chosen to prevent the degradation of storage lipids and the subsequent generation of acetyl-CoA for β-oxidation pathway, which could confound the interpretation of flux through our engineered pathways. To compare the performance of the novel pathways, we constructed three different strains (**Fig. 2B**) by integrating either ACC1 (canonical pathway), the ILM pathway (IboH-LtaE-MCR-C, using L-glutamate as substrate, **Fig. 1A**), or the AMM pathway (ADC-MAO-MCR-C, using L-aspartate as substrate, **Fig. 1B**) into the *DGA1* locus under well-defined promoters (Fig. 2B). The DGA1 gene, encoding a diacylglycerol acyltransferase essential for triglyceride biosynthesis, was chosen as the genome integrated loci to divert malonyl-CoA flux away from lipid synthesis and toward polyketide production.

**Fig. 2.**
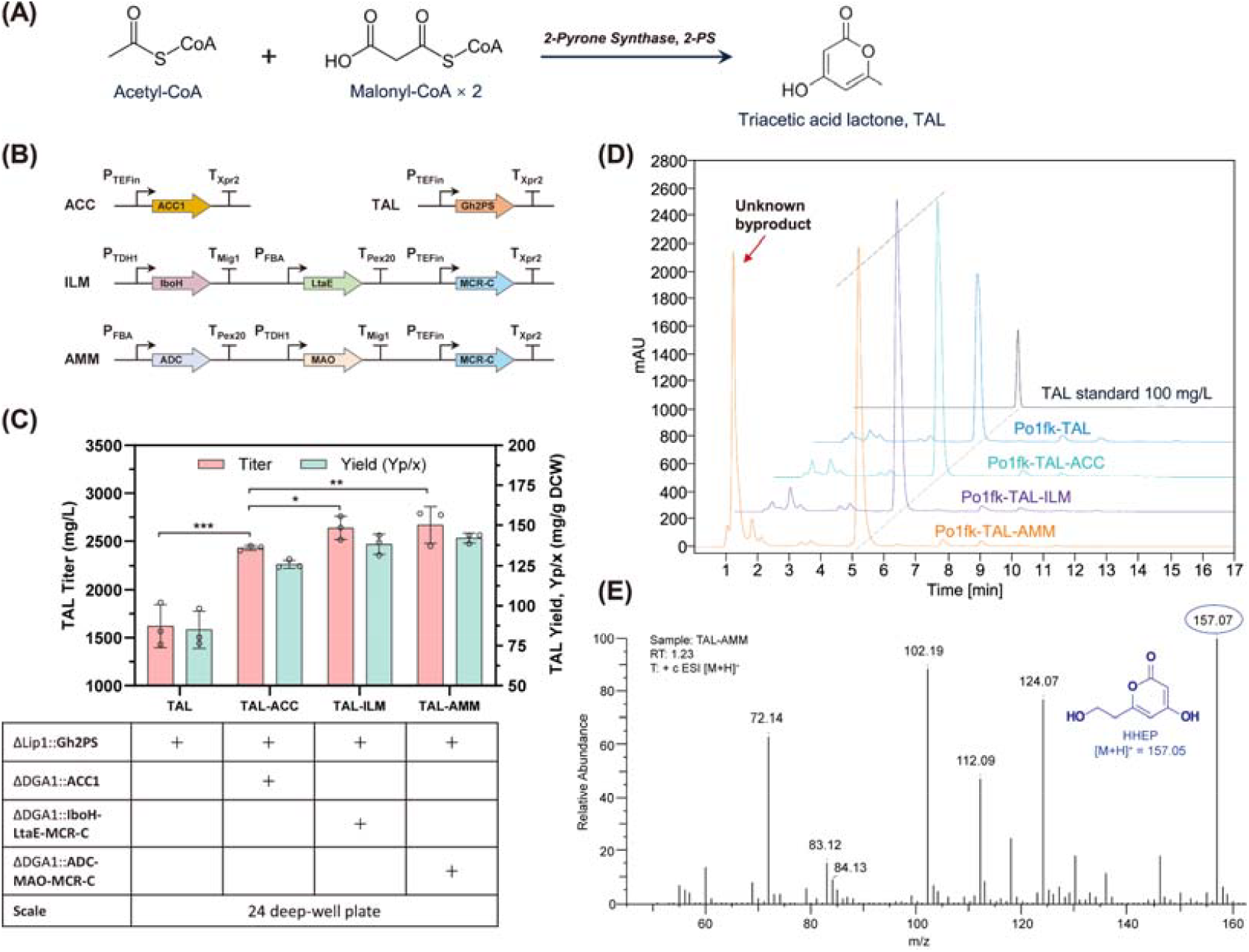
Testing the functionality of the novel malonyl-CoA pathways. (**A**) The simplest polyketide pathway encoded by 2-pyrone synthase (2-PS) was used as a reporter molecule; (B) Three TAL pathways, TAL-ACC, TAL-ILM, and TAL-AMM tested in this work; (**C**) TAL titer and specific TAL yield from the engineered strains in deep-well plates using YPD80 medium. Experiments were performed in triplicate, and the data is reported as mean ± SD. *** indicates statistically highly significant (*p* < 0.001), ** indicates statistically significant (*p < 0*.01), * indicates statistically significant (*p < 0*.05); (D) HPLC profile of TAL and an unknown by-product from the engineered strains; (**E**) LC-MS spectra of the unknown metabolite from the TAL-AMM strain.

TAL production in deep-well plate cultures (**Fig. 2C**) indicates that both ILM and AMM pathways outperformed the canonical ACC1 route in terms of titer and specific yield (mgTAL/gDCW). Specifically, the strain carrying the ILM pathway achieved a TAL titer of 2638.8 mg/L, while the AMM pathway led to a slightly higher TAL titer of 2669.6 mg/L (**Fig. 2C**), both exceeding the ACC1 control (2431.1 mg/L). This suggests that the oxidative deamination and decarboxylation steps in the novel pathways (**Fig. 1**) effectively converted the amino acid precursors into malonyl-CoA, overcoming the inherent limitations of ATP-dependent carboxylation in the ACC1 pathway (**Table 1**). The higher TAL yield observed in the AMM pathway could be attributed to a more favorable kinetic profile of the ADC-MAO cascade (**Table 2**).

Interestingly, HPLC analysis revealed an unknown by-product in the strain carrying the AMM pathway (**Fig. 2D**). This unknown peak persists in the shaking flask culture (supplementary **Fig. S1B**). Moreover, in shaking flasks, AMM strain produced less TAL than the strain carrying ILM pathway (supplementary **Fig. S1A**), possibly due to competition from an unknown by-product that accumulates in the AMM pathway (supplementary **Fig. S1B**). This byproduct is likely derived from the β-alanine or MalSA intermediates. We next identified the structure of the unknown metabolite (**Fig. 2D**) observed in the AMM strain. LC–MS analysis (**Fig. 2E**) revealed a major species eluting at 1.26 min (supplementary **Fig. S2**), with a molecular weight of 157 Dalton and characteristic fragmentation ions at *m/z* = 124, 112, 102, 84 and 72 under positive (M+1) ionization modes. By comparing these fragmentation patterns against standard databases and predicted mass spectra (supplementary **Fig. S3**), we characterized this molecule as 4-hydroxy-6-hydroxyethyl-2-pyrone (HHEP) with a chemical formula C_7_H_8_O_4_.

### 2.3 3-HP-CoA serves as the priming unit for the nonnatural polyketide HHEP

Previously, the β-alanine pathway has been identified as the most economically attractive route to synthesize 3-hydroxypropionate (3-HP) (*32*). Inspection of TAL biosynthetic mechanism (**Fig. 3A**) led us to hypothesize that HHEP was generated as a polyketide by-product when 3-hydroxypropanoyl-CoA (3-HP-CoA) was used as an alternative priming molecule for 2-pyrone synthase (2PS) (**Fig. 3B** and **3C**) (*33*). In the AMM strain, β-alanine generated from aspartate decarboxylase (ADC) can be oxidized to form MalSA, which further is reduced to 3-HP by host enzyme HBD1 encoding 3-hydroxyacyl-CoA dehydrogenase (**Fig. 3B** lower panel). In fact, 3-HP-CoA is also the degradation product of methionine, threonine, leucine, and valine (**Fig. 3A** upper panel) from the media containing YPD and peptone. When we blasted the genome of *Y. lipolytica*, we identified that at least one gene (YALI0D06325) encodes an enzyme highly conserved with BAPAT encoding alanine aminotransferase which converts β-alanine to MalSA (supplementary **Fig. S6**); and likewise, YALI0C08811 and YALI0F05962/YALI0D06215g are highly conserved with HBD1 encoding 3-hydroxyacyl-CoA dehydrogenase, and ACS2/EHD3 encoding 3-hydroxypropanoyl-CoA synthetase (supplementary **Fig. S7-S9**), respectively. Theoretically, when 2PS encounters 3-HP-CoA as a priming unit, it could condense with two molecules of malonyl-CoA to form a hydroxymethyl triketide intermediate (**Fig. 3C**). This intermediate, undergoes a series of intramolecular nucleophilic substitutions and spontaneous lactonization, ultimately cyclizes and rearranges to yield the more stable lactone 4-hydroxy-6-hydroxyethyl-2-pyrone (HHEP) (**Fig. 3E**) following a similar pattern as 4-hydroxy-6-methyl-2-pyrone (TAL) (**Fig. 3D**) does. Interestingly, we found HHEP was synthesized in all the strains expressing 2PS (**Fig. 2D**, supplementary **Fig. S4** for ILM-TAL strain and supplementary **Fig. S5** for TAL strain), albeit HHEP was mostly accumulated in the AMM strain (supplementary **Fig. S1B** and **Fig. S2**). The non-exclusive production of HHEP from all the tested strains suggests that the primer unit, 3-HP-CoA, is derived at low levels from the degradation of branched-chain amino acids (e.g., valine, methionine, threonine) (**Fig. 3B** upper panel) or generated from the reduction of MalSA via β-alanine metabolism (**Fig. 3B** lower panel).

**Fig. 3.**
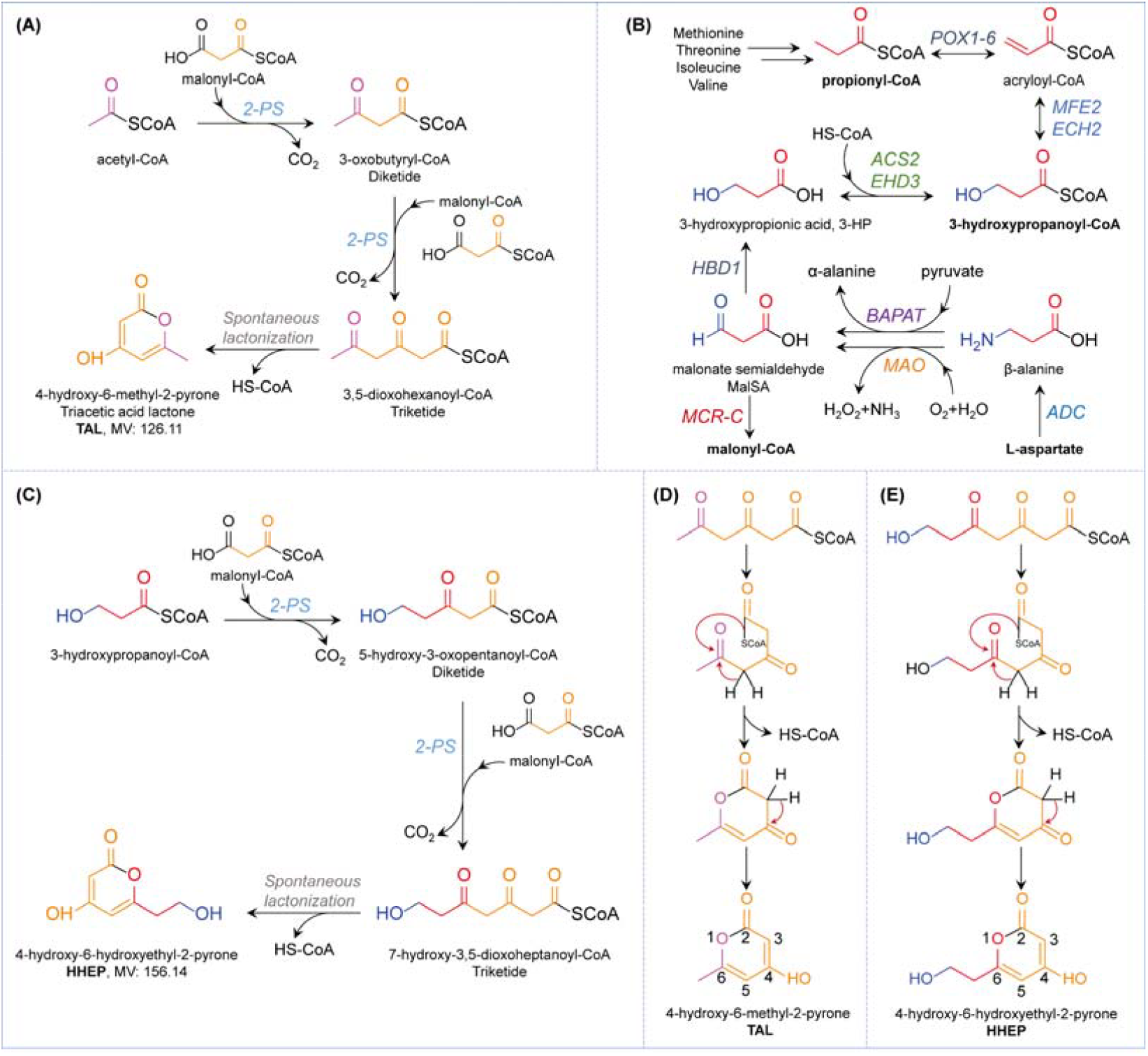
Deciphering the biosynthetic route of the nonnatural polyketide HHEP. (**A**) Biosynthetic route of TAL by 2PS. (**B**) Generation of 3-HP and 3-HP-CoA from endogenous amino acid metabolism (upper panel) and β-alanine from L-aspartate decarboxylation (lower panel). BAPAT: β-alanine-pyruvate aminotransferase; MAO: monoamine oxidase; HBD1: 3-hydroxyacyl-CoA dehydrogenase; ACS2/EHD3: 3-hydroxypropanoyl-CoA synthetase; POX1-6: Fatty-acyl-CoA oxidase; MFE2: Multifunctional β-oxidation enzyme, hydratase/dehydrogenase; ECH2: Enoyl-CoA hydratase. (**C**) 3-HP-CoA serves as a priming molecule to synthesize HHEP by 2PS. Spontaneous lactonization of triketide intermediates leads to the formation of triacetic acid lactone (**D**) and 4-hydroxy-6-hydroxyethyl-2-pyrone (**E**).

To validate that HHEP is derived from 3-HP, we next fed 34 mM β-alanine to the TAL-ACC strain and observed a substantially increased level of 3-HP in shaking flask (Fig. **4A** and supplementary **Fig. S10A**). The presence of 3-HP was also confirmed by LC-MS analysis (**Fig. 4B**). For example, TAL-ACC strain produced 3042.0 mg/L of 3-HP with feeding of 34 mM β-alanine, more than 3-fold higher than the same strain without feeding of β-alanine (831.8 mg/L) (**Fig. 4C** and supplementary **Fig. S10A**). These facts confirm that endogenous BAPAT (YALI0D06325) and HBD1 (YALI0C08811) can convert β-alanine to 3-HP (**Fig. 3B**). To directly validate that HHEP is derived from 3-HP-CoA, we next fed TAL-AMM with 34 mM of 3-HP. However, the addition of 34 mM 3-HP was found to inhibit cell growth, likely due to its inherent toxicity (supplementary **Fig. S11A**). Therefore, we reduced the 3-HP supplementation to 10 mM, while simultaneously supplying 34 mM L-aspartate to ensure sufficient malonyl-CoA availability for the condensation of 3-HP-CoA and malonyl-CoA into HHEP. The results demonstrated that feeding 10 mM 3-HP significantly increased the HHEP titer to 6359.1 mg/L, which is 22.5% higher than that of the non-fed group (5191.3 mg/L) (**Fig. 4D** and supplementary **Fig. S11C**). These findings confirm that the endogenous 3-hydroxypropanoyl-CoA synthetase can activate 3-HP to 3-HP-CoA for the synthesis of the polyketide HHEP (**Fig. 3B**). When only ADC was expressed, the TAL-A strain produced basal level of TAL (751.0 mg/L) and HHPE (542.4 mg/L) (**Fig. 4E** and supplementary **Fig. S10B**). However, when the entire AMM pathway was expressed, the TAL-AMM strain produced 6-fold more HHPE (3415.1 mg/L) compared to the TAL-A strain (**Fig. 4E** and Supplementary **Fig. S10C**). This result showcases the critical roles of monoamine oxidase (MAO) and MCR to boost intracellular malonyl-CoA and improve HHEP synthesis (**Fig. 3B** and **Fig. 3C**). The discrepancy of TAL and HHEP titer enhancement indicates the intracellular scarcity of acetyl-CoA relative to 3-hydroxypropanoyl-CoA in the TAL-AMM strain, alternatively, 2PS may prefer 3-HP-CoA over acetyl-CoA as the priming molecule. Taken together, these results confirm that HHEP arises as a nonnatural polyketide by-product when 3-HP-CoA participates in the 2-PS condensation reaction (**Fig. 3B** and **Fig. 3C**).

**Fig. 4.**
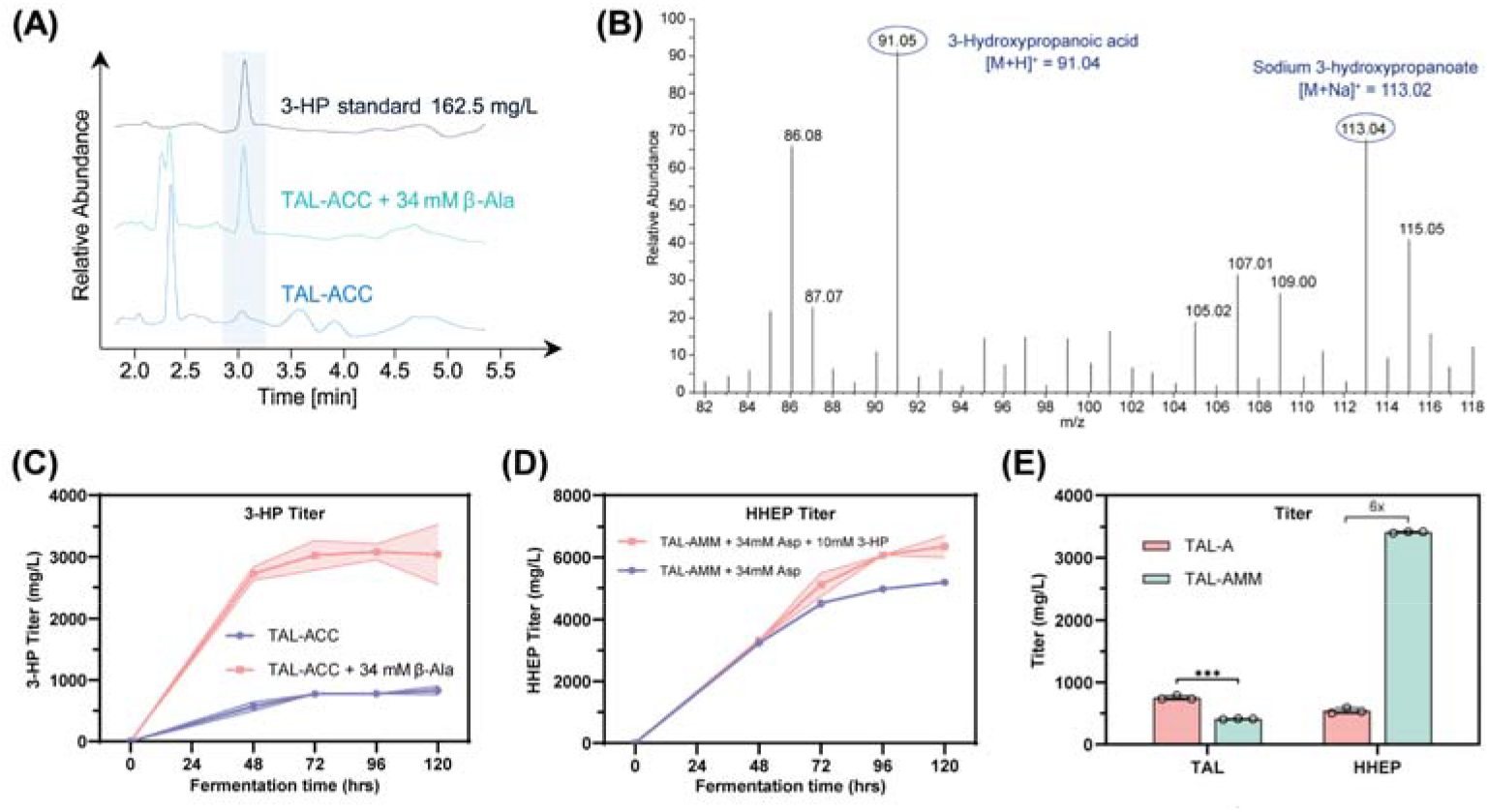
Validating 3-HP as an intermediate for HHEP synthesis. (**A**) HPLC profile of 3-HP in the TAL-ACC strain with feeding of 34 mM β-alanine; (**B**) LC-MS characterization of 3-HP in the TAL-ACC strain; (**C**) Production of 3-HP in the TAL-ACC strain with or without feeding of 34 mM β-alanine in shaking flask using YPD80 medium. β-Ala feeding: 5 mM, 5 mM, 8 mM, 8 mM, and 8 mM were added at 24 h, 36 h, 48 h, 60 h, and 72 h, respectively, resulting in a total supplement of 34 mM; (**D**) Production of HHEP in the TAL-AMM strain with or without feeding of 10 mM 3-HP in shaking flask using YPD80 medium. 3-HP feeding: 1.4 mM, 1.4 mM, 2.4 mM, 2.4 mM, and 2.4 mM were added at 24 h, 36 h, 48 h, 60 h, and 72 h, respectively, resulting in a total supplement of 10 mM. L-Aspartate feeding: 5 mM, 5 mM, 8 mM, 8 mM, and 8 mM were added at 24 h, 36 h, 48 h, 60 h, and 72 h, respectively, resulting in a total supplement of 34 mM; (**E**) TAL and HHEP production in the TAL-A and TAL-AMM strain after 48 hours of fermentation in deep-well plates using YPD80 medium. TAL-A is the strain with expression of 2PS and ADC. TAL-AMM is the strain with expression of 2PS and the entire AMM pathway.

HHEP is a structural analog of the natural product Kojic acid, which is a well-known whitening and skincare natural product. This is the first report on the synthesis of the nonnatural polyketide HHEP, and the biological activity of HHEP has yet to be discovered. Furthermore, HHEP holds promise as a versatile bio-based platform compound. It could serve as a precursor for the synthesis of functional polymers, including biodegradable polyesters and UV-curable resins, through esterification or oxidative modification. In addition, the conjugated system may enable its application as an antioxidant, UV-absorbing, or cross-linkable additive in coatings and materials. These properties suggest that HHEP represents a valuable compound for developing sustainable, high-value chemicals and polymeric materials.

### 2.4 Feeding amino acids to boost polyketide and lipid production in shaking flask

We next tested whether feeding L-glutamate or L-aspartate to the ILM and AMM strains will boost malonyl-CoA derivatives production, including TAL, HHEP and lipid. Since our designed pathway is oxygen-dependent, we cultured the ILM strain in shaking flask to improve oxygen supply. After feeding 34 mM L-glutamate to the TAL-ILM strain, both TAL and HHEP titer were increased in the shaking flasks (**Fig. 5A**). TAL yield was increased by 25% compared to the same strain without L-glutamate feeding (supplementary **Fig. S12A** and **Fig. S12D**). This result indicated that feeding with L-glutamate boosted malonyl-CoA pools and polyketide synthesis in the TAL-ILM strain. HHEP instead decreased after 72 hours (supplementary **Fig. S12B**), indicating the possible degradation of HHEP by the host.

**Fig. 5.**
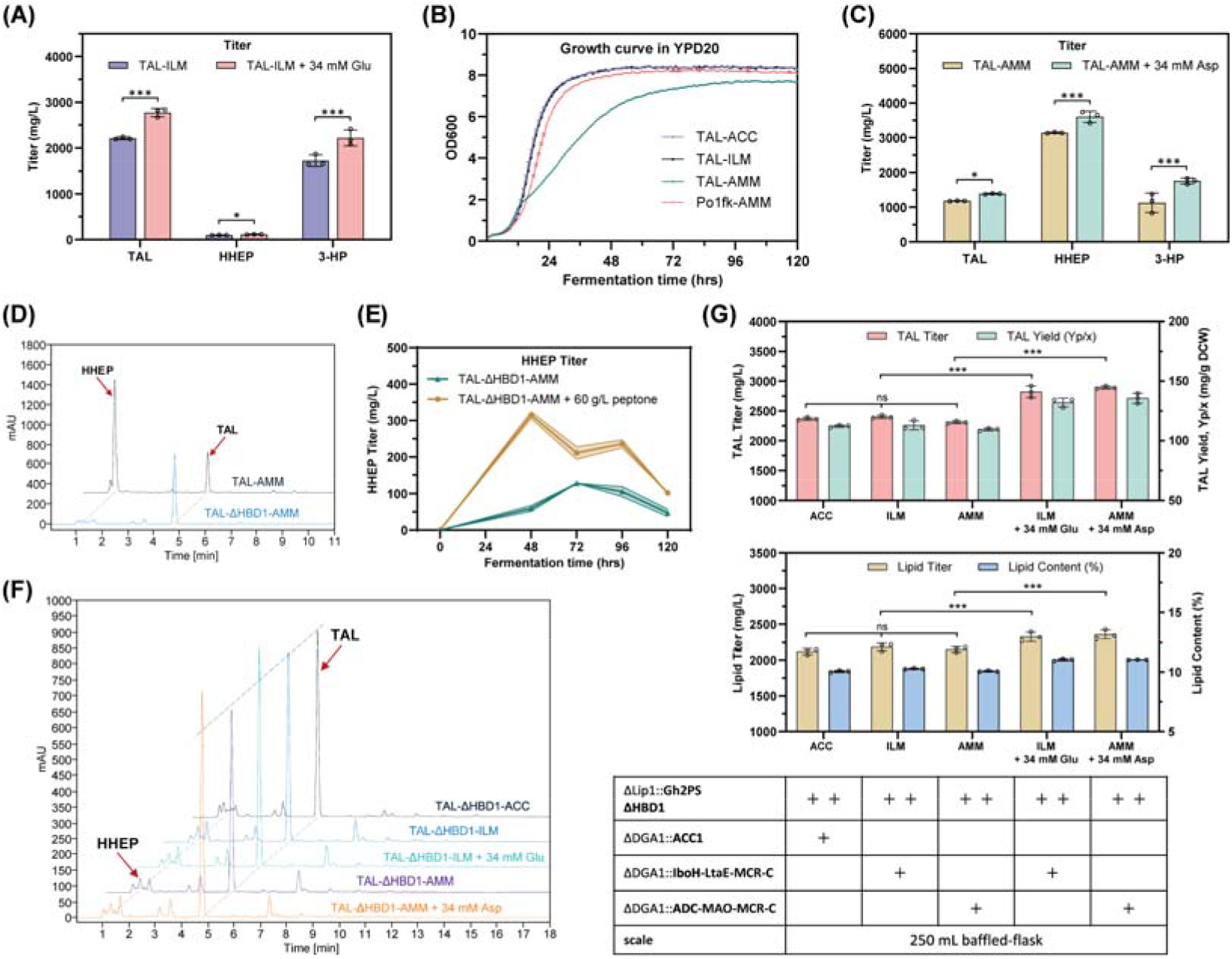
Feeding L-glutamate or L-aspartate to improve polyketide production in TAL-ILM or TAL-AMM strain. (**A**) TAL, HHEP and 3-HP production in TAL-ILM strain with or without feeding 34 mM L-glutamate in 250 mL baffled-flask using YPD80 medium. L-Glutamate feeding: 5 mM, 5 mM, 8 mM, 8 mM, and 8 mM were added at 24 h, 36 h, 48 h, 60 h, and 72 h, respectively, resulting in a total supplement of 34 mM; (**B**) Growth fitness of HHEP-producing strain (TAL-AMM) is reduced compared to other TAL-producing strain; (**C**) TAL, HHEP and 3-HP production in TAL-AMM strain with or without feeding of 34 mM L-aspartate in 250 mL baffled-flask using YPD80 medium. L-Aspartate feeding: 5 mM, 5 mM, 8 mM, 8 mM, and 8 mM were added at 24 h, 36 h, 48 h, 60 h, and 72 h, respectively, resulting in a total supplement of 34 mM; (**D**) HPLC profile show that the HHEP peak returned to basal level almost undetectable when HBD1 gene was deleted; (**E**) Production of HHEP in the TAL-ΔHBD1-AMM strain with or without feeding of 60 g/L peptone in shaking flask using YPD80 medium. Peptone feeding: 10 g/L, 10 g/L, 10 g/L, 15 g/L, and 15 g/L were added at 24 h, 36 h, 48 h, 60 h, and 72 h, respectively, resulting in a total supplement of 60 g/L; (**F**) HPLC profile of TAL and HHEP from the engineered strains after knocking out HBD1: the product shifts to TAL accumulation; (**G**) TAL titer, specific TAL yield, lipid titer and lipid content from the engineered strains with or without amino acid feeding in 250 mL baffled-flask using YPD80 medium. L-Glutamate or L-aspartate feeding: 5 mM, 5 mM, 8 mM, 8 mM, and 8 mM were added at 24 h, 36 h, 48 h, 60 h, and 72 h, respectively, resulting in a total supplement of 34 mM. Experiments were performed in triplicate, and the data is reported as mean ± SD. *** indicates statistically highly significant (p < 0.001), * indicates statistically significant (*p < 0*.05), ns indicates statistically significant (p > 0.05).

While cell growth in the TAL-ILM, TAL-ACC and AMM strains remain unchanged; the TAL-AMM strain exhibits pronounced growth-retardation (**Fig. 5B**), possibly due to the inhibitory effect of HHEP. Using TAL as an internal standard, we determined that TAL-AMM strain produced 3140.9 mg/L of HHEP (**Fig. 5C**). After feeding 34 mM L-aspartate to TAL-AMM strain, the production of TAL, HHEP and 3-HP increased by 17.4%, 14.7% and 55.5%, respectively (**Fig. 5C** and supplementary **Fig. S13B**). And the total polyketide (both TAL and HHEP) was boosted to 5.0 g/L (**Fig. 5C**), which is more than 1.9-fold of the total polyketides (TAL and HHEP) than the ACC pathway (supplementary **Fig. S1A**).

We next deleted HBD1, to eliminate interference from the endogenous 3-HP-CoA-related pathways (**Fig. 3B** and **3C**). Compared to TAL-AMM strain, the TAL-ΔHBD1-AMM strain exhibited significantly lower HHEP production (**Fig. 5D**). However, HHEP was still detectable in all the HBD1 knockout strain (**Fig. 5D**), suggesting that the precursor of HHEP, 3-hydroxypropanoyl-CoA, may also be derived from the endogenous branched-chain amino acid degradation (**Fig. 3B** upper panel). Therefore, we next fed the TAL-ΔHBD1-AMM strain with 60 g/L peptone to provide abundant branched-chain amino acids. As a result, the HHEP titer increased by 1.5-fold compared with cultures grown on YPD medium alone (**Fig. 5E** and supplementary **Fig. S14A**). This indicates that 3-HP-CoA derived from branched-chain amino acid catabolism can contribute to the baseline HHEP production across all 2PS-expressing strains. Cell growth fitness analysis (supplementary **Fig. S14B**) further revealed that deletion of HBD1 restored normal growth in the TAL-AMM strain, indicating that HHEP indeed acts as a growth-inhibitory compound. Taken together, removal of HBD1 has allowed for a more straightforward assessment of the pathway’s performance by eliminating a major malonyl-CoA competing pathway.

To assess the efficiency of different malonyl-CoA pathways, we next constructed three engineered strains based on HBD1-deleted chassis, the TAL-ΔHBD1-ACC strain produced 2368.9 mg/L TAL from glucose, while the ILM and AMM strains yielded 2401.4 mg/L and 2310.4 mg/L, respectively. Similarly, the TAL-ΔHBD1-ACC strain produced 2121.9 mg/L lipid from glucose, while the ILM and AMM strains yielded 2187.1 mg/L and 2152.0 mg/L, respectively. These results indicate that the ILM and AMM pathways provide malonyl-CoA at a level comparable to that of ACC pathway (**Fig. 5G**). We next fed 34 mM L-glutamate to the ILM strain, TAL production was increased from 2401.4 mg/L to 2822.6 mg/L, and lipid production was increased from 2187.1 mg/L to 2329.0 mg/L. Similarly, feeding 34 mM L-aspartate to the AMM strain raised TAL titer from 2310.4 mg/L to 2893.4 mg/L and lipid titer from 2152.0 mg/L to 2356.9 mg/L, representing increases of 25.2% and 9.5%, respectively. (**Fig. 5F** and **Fig. 5G**). These results confirm that the new amino acid-derived malonyl-CoA pathways effectively boost polyketide and lipid production in *Y. lipolytica*.

Interestingly, significantly elevated HHEP titers were observed specifically in the TAL-AMM strain (**Fig. 4D**), which may result from the MalSA precursor overflow effect. The TAL-AMM strain generates malonate semialdehyde (MalSA) from two primary sources: engineered MAO and endogenous BAPAT steps (**Fig. 3B** lower panel). Although MCR step is designed to oxidize MalSA to malonyl-CoA, the endogenous HBD1 effectively competes with MCRA pool, reducing MalSA to 3-HP, which is subsequently activated to 3-HP-CoA by EHD3 (Fig. 3B). This creates an abundant supply of the starting unit 3-HP-CoA for 2PS, leading to high-level HHEP synthesis. By deleting HBD1 in the TAL-AMM strain, HHEP production reduced to a level comparable to other 2PS-expressing strains, which 3-HP-CoA can only be derived from the degradation of the branched-chain amino acids in the media (**Fig. 3B** upper panel). In contrast, the TAL-ILM strain only produces MalSA from the aldolase LtaE (**Fig. 1B**), resulting in less 3-HP-CoA formation and consequently lower HHEP titer.

### 2.5 Feeding L-aspartate to boost polyketide and lipid production in bioreactor

To evaluate the effect of L-aspartate feeding on polyketide production at a larger scale, we conducted a 1 L bioreactor fermentation using the TAL-AMM strain, with or without the addition of 100 mM L-aspartate. The combined titer of TAL and HHEP in the L-aspartate-fed culture reached a peak of 6955.2 mg/L at 108 h, which is a 26.4% increase compared to the maximum titer of 5500.5 mg/L in the control strain without L-aspartate feeding (**Fig. 6A**). This substantial improvement suggests that L-aspartate feeding enhances malonyl-CoA supply toward total polyketide biosynthesis. With L-Asp feeding, HHEP reached a peak of 5368.6 mg/L, representing a 49.2% increase compared to the control group (3597.3 mg/L) (**Fig. 6B**). Notably, HHEP reached 5113.0 mg/L by 60 hours of fermentation, when only 70 g/L of glucose was consumed (**Fig. 6F**). HHEP accumulation slows down after 60 hours, despite continued glucose consumption. This may indicate degradation of the product HHEP.

**Fig. 6.**
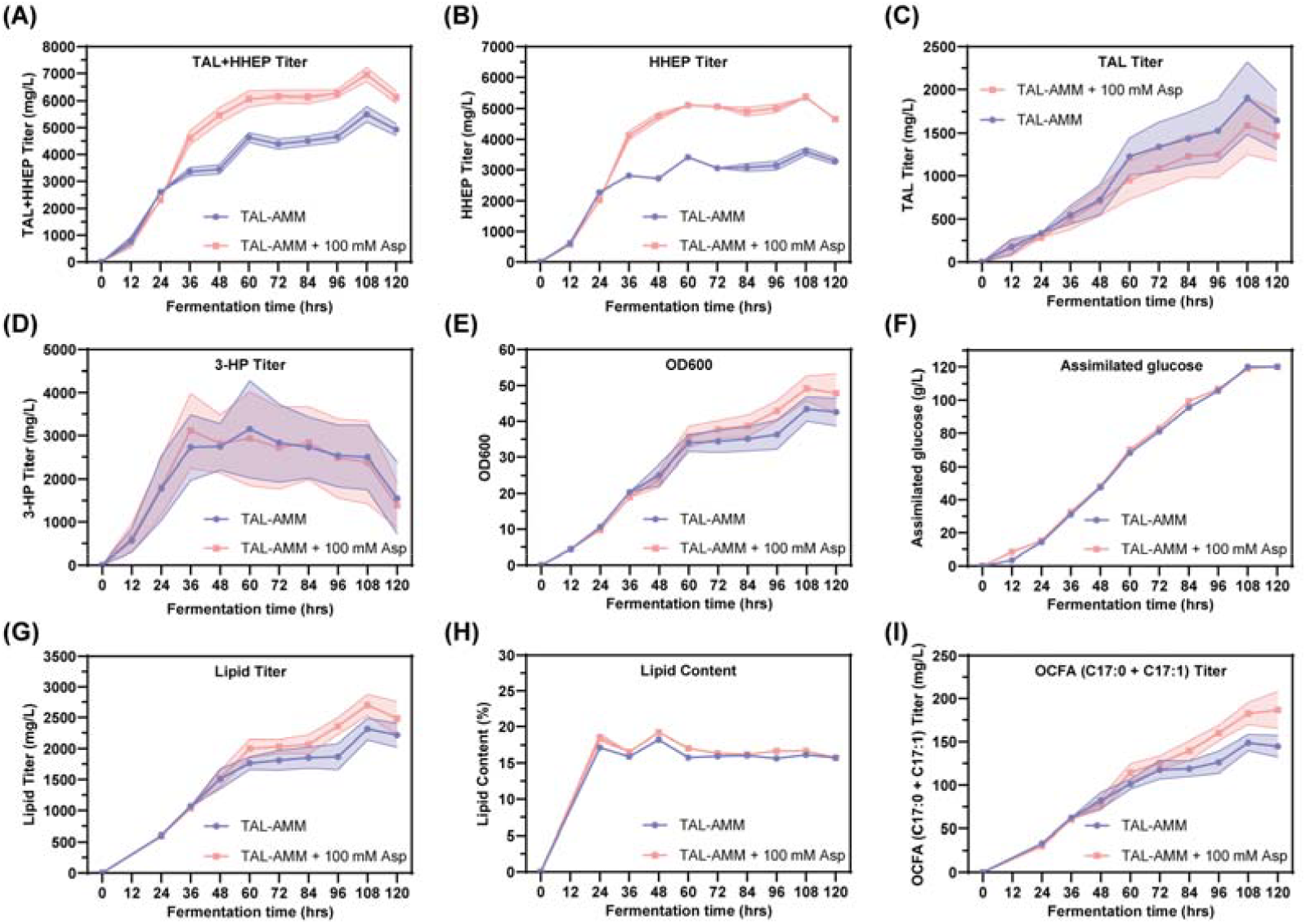
Feeding L-aspartate to improve polyketide and lipid production in TAL-AMM strain in a 1.0 L mini-bioreactor. TAL+HHEP(**A**), HHEP(**B**), TAL(**C**) and 3-HP(**D**) production in TAL-AMM strain with or without feeding of 100 mM L-aspartate in a 1L bioreactor. L-Aspartate feeding: 5 mM, 10 mM, 15 mM, 15 mM, 15 mM, 20 mM, and 20 mM were added at 0 h, 12 h, 24 h, 36 h, 48 h, 60 h, and 72 h, respectively, resulting in a total supplement of 100 mM; (**E**) Optical density OD600 during fermentation in a 1L bioreactor; (**F**) Amount of glucose consumed during fermentation in a 1L bioreactor; Total Lipid titer (**G**), lipid content (**H**) and total odd-chain fatty acid (OCFA) production in TAL-AMM strain with or without feeding of 100 mM L-aspartate in a 1L bioreactor.

Interestingly, TAL production decreased in the L-aspartate-fed culture compared to the control, particularly after 48 hours (**Fig. 6C**). This suggests that with excess L-aspartate, a greater portion of the precursor malonyl-CoA is channeled toward HHEP biosynthesis rather than TAL accumulation. The increased availability of L-aspartate likely facilitates the formation of the HHEP precursor 3-hydroxypropionyl-CoA (3-HP-CoA), thereby leading to a substantial increase in HHEP. This shift in product distribution reinforces the role of L-aspartate as a flux-enhancing agent that drives carbon flowing through the AMM pathway. Furthermore, the 3-HP titer in the L-aspartate-fed culture peaked at 3119.0 mg/L, equivalent to 3152.2 mg/L maximum in the control strain. Notably, 3-HP production gradually decreased after 60 hours, regardless of whether L-aspartate was fed (**Fig. 6D**). This suggests a shift in metabolic flux from 3-HP to HHEP: 3-HP was converted to 3-HP-CoA promoting an increased flux toward HHEP.

Cell growth slightly improved with L-aspartate supplementation, reaching a final OD600 of 49.2, compared to 43.4 in the control (**Fig. 6E**). Despite differences in product profiles, glucose consumption was identical between both conditions, with a total consumption of 120 g/L at the end of fermentation (**Fig. 6F**). This indicates that the observed differences in product titers were not due to changes in glucose uptake but rather due to increased supply of malonyl-CoA through the amino-acid-derived malonyl-CoA pathways.

For lipid titer, L-aspartate supplementation led to a consistent increase in total lipid titer throughout the fermentation process (**Fig. 6G**). The maximum lipid titer of the L-Asp supplemented culture reached 2703.7 mg/L, representing a 17.0% increase over the control (2311.6 mg/L). The lipid content (% of cell dry weight) exhibited only a slight increase under L-aspartate feeding, peaking at 19.2% (**Fig. 6H**). This indicates that L-aspartate raises the absolute lipid titer by promoting malonyl-CoA synthesis. In addition, we observed a pronounced increase in odd-chain fatty acid (OCFA; C17:0 and C17:1) production following L-aspartate supplementation. The maximum OCFA titer in the supplemented culture reached 186.5 mg/L, compared to 148.9 mg/L in the control, corresponding to a 25.3% increase (**Fig. 6I**, supplementary **Fig. S15A**). This enhancement can be attributed to the ability of L-aspartate to boost the supply of propionyl-CoA, the starting unit for OCFA biosynthesis (**Fig. 3B**). OCFA synthesis also needs malonyl-CoA as the extender units. This finding is consistent with earlier observations that L-aspartate feeding redirects metabolic flux toward the synthesis of polyketide and lipid within the AMM pathway.

Together, these results demonstrate that L-aspartate feeding is an effective strategy to enhance high-value polyketide production in engineered *Y. lipolytica*. Both L-aspartate and L-glutamate are among the most abundant amino acids in food and animal farming waste. Different from the reported transaminase-MCR route (*20*), our re-designed malonyl-CoA pathway leverages hydroxylation (in the case ILM) or oxidative deamination (in the case AMM) with oxygen consumption, and therefore it provides a large driving force to compensate the relatively inefficient MCR step. When coupled together, the novel ILM and AMM pathway efficiently converted L-glutamate or L-aspartate to malonyl-CoA and improved the production of both TAL and HHEP. These novel pathways are particularly useful to produce malonyl-CoA-derived compounds under aerobic conditions, potentially benefiting large-scale fermentations. Our results established a promising and scalable polyketide production platform from low-cost amino acid precursors or waste protein streams.

## 3. Conclusions

Herein we have designed and validated two malonyl-CoA biosynthetic pathways in *Y. lipolytica*, utilizing L-glutamate and L-aspartate as substrates. These pathways provide an efficient and ATP-independent alternative for metabolic engineering applications, which expands nature’s capacity to produce malonyl-CoA-derived chemicals in *Y. lipolytica*. By leveraging oxidative deamination and decarboxylation reactions, both pathways effectively redirected metabolic flux toward malonyl-CoA, significantly enhancing the biosynthesis of triacetic acid lactone (TAL), a new-to-nature polyketide HHEP, lipid and odd chain-length fatty acids. The strain with AMM pathway produced a novel polyketide HHEP, at 6.4 g/L, highlighting the utility of the amino acid-derived malonyl-CoA pathway and the superior metabolic versatility of *Y. lipolytica*. The total polyketide production reached 7.0 g/L in a 1L bioreactor. Our findings demonstrated that low-cost amino acids or protein waste streams could be used as efficient raw materials to manufacture high-value polyketides. The study also reveals thermodynamic coupling could be used to overcome inefficient enzymatic steps. Overall, this work represents an important step toward sustainable and cost-effective production of polyketides by rewiring the amino acid metabolism in microbes.

## 4. Materials and methods

### 4.1 Strains and cultural conditions

All engineered *Y. lipolytica* strains in this research were derived from Po1fk, a commonly used laboratory strain of *Y. lipolytica* that is leucine- and uracil-prototrophic (Po1g-Δura3-Δku70::loxP). *Escherichia coli* DH5α was cultured in LB medium supplemented with 100 mg/mL ampicillin for plasmid construction and extraction. *Y. lipolytica* strains were cultured in YPD medium for genomic DNA extraction or linearized plasmid transformation. Engineered strains were fermented in YPD-based medium to produce triacetic acid lactone. SD-Ura and SD-Leu plates were used to isolate positive *Y. lipolytica* transformants. The Cre-loxP system was used to eliminate the URA3 selection marker (*34*). To conduct 24-deep-well plates or 250 mL shake-flask fermentation, the strain was grown in 3 mL YPD medium for 1 day to prepare a seed culture. A portion of the seed culture was utilized to inoculate 3.5 mL or 15 mL of YPD medium with 80 g/L glucose and additional glutamic acid or aspartic acid, with the aim of achieving an initial OD600 value of 0.2, followed by fermentation at 30°C and 250 rpm for 120 h. All engineered strains utilized in fermentation were prototrophic. The formulations of LB medium, YPD medium, SD-Ura plate, SD-Leu plate, and YPD-5-FOA plate were outlined in the supplementary **materials**.

### 4.2 Genetic engineering and strain construction

In this research, expression cassette plasmids for individual genes were generated by Gibson assembly and verified by DNA sequencing. Subsequently, we assembled multiple expression cassettes on an integrative vector using a subcloning method of digestion and ligation, following YaliBrick assembly protocols (*35, 36*). All plasmids constructed are listed in the supplementary **Table S1**. The plasmid (*35*) used for gene integration consists of two 1∼1.5 kb homology arms, which correspond to sequences 5’-upstream and 3’-downstream of the integration site, and a loxP-URA3-loxP cassette, the latter of which was used to recover the URA3 screening marker by using Cre-loxP recombination system. The plasmid used for gene integration was inserted into the *Y. lipolytica* chromosome for gene overexpression. Homology arms of integration sites, as well as terminators and promoters, were amplified by PCR using the genomic DNA from strain Po1fk or other constructed plasmids as templates. The exogenous gene coding sequences were codon-optimized and synthesized by Genewiz (Suzhou, China). *Y. lipolytica* endogenous gene *Yl*ACC1 (YALI0C11407g) devoid of intron sequence was amplified through PCR utilizing the genomic DNA of strain Po1fk as the template. The gene sequences that have been codon optimized are provided in the supplementary **Table S2**. All cloning primers and check primers are listed in supplementary **Table S3**.

### 4.3 Y. lipolytica *transformation and strain verifications*

The strain to be transformed was cultured in YPD medium for 24 hours, after which the yeast cells were collected and resuspended in PBS buffer to wash away the YPD medium. Subsequently, 50% PEG4000, 2 M lithium acetate, salmon sperm DNA, and a linearized plasmid were added for homologous recombination transformation (*37*). Positive clones were selected on CSM-Ura plates, and genomic DNA from the clones was extracted using the Lysis Buffer with Microorganism Direct PCR Kit (Takara, Beijing) for PCR-based identification. A single positive clone identified by PCR was then inoculated in YPD medium for 24 hours and transformed with the pYLXP’-Cre plasmid to eliminate the URA3 marker gene. The resulting colonies were streaked on both YPD-5-FOA and CSM-Ura plates. Colonies that grew on YPD-5-FOA plates but failed to grow on CSM-Ura plates were selected for PCR analysis to confirm the successful elimination of the URA3 marker gene. Strains lacking the URA3 marker gene were then subjected to another round of linearized plasmid transformation and screening. All strains constructed in this work are listed in supplementary **Table S4**.

### 4.4 Dry cell weight quantification and triacetic acid lactone extraction

The cell growth was monitored by measuring the optical density at 600 nm (OD600) with a UV–vis spectrophotometer that could also be converted to dry cell weight (DCW) according to the calibration curve (DCW: OD600 = 0.33:1 (g/L). The fermentation broth was centrifuged at 12,000 rpm for 3 min, and the culture was used for analyzing the concentration of TAL. The reported titers for TAL and HHEP represent the total (intracellular and extracellular) concentrations. Metabolite extraction was performed by subjecting the entire culture broth, including cells, to freeze-thaw cycles and mechanical grinding to ensure complete release of intracellular products prior to analysis. In general, the extracellular TAL or HHEP accounts for more than 85% of the production. For complete extraction of TAL, 100 μL fermentation culture was resuspended in 700 μL of ultrapure water containing plastic particles as abrasives. The cell culture was grinded using a cryo-grinding machine at -20 °C for 20 cycles of 40 s each, followed by centrifugation at 12,000 rpm for 3 min. After the mixture was centrifuged, the supernatant was taken for high performance liquid chromatography (HPLC) analysis.

### 4.5 Characterization and quantification of TAL, 3-HP, and unknown peak

An Agilent 1260 Infinity II instrument was utilized for HPLC analysis, with a variable wavelength detector (VWD) and a C18 column (4.6 mm × 100 mm). The HPLC analysis was carried out using a mobile phase consisting of solvent A (0.1% formic acid in water) and solvent B (methanol). A gradient elution was applied starting with 10% B, increasing to 100% B over 20 minutes, followed by 3 minutes at 100% B. The flow rate was 1.0 mL/min, with a column temperature of 30°C. Detection was performed at 280 nm, and the injection volume was 10 µL (*38*). Retention time of triacetic acid lactone is about 5.2 min. The HPLC analysis for 3-hydroxypropionate (3-HP) was carried out using a single mobile phase (0.1% formic acid in water - methanol (95:5, v/v) mixture), and the total elution time was 10 minutes. The flow rate was 1.0 mL/min, with a column temperature of 30°C. Detection was performed at 210 nm for 3-HP, and the injection volume was 10 µL. Retention time of 3-HP is about 3.0 min. TAL and 3-HP standard was purchased from Macklin, China.

The unknown compound was characterized by using a Thermo Scientific Q Extractive mass spectrometer equipped with an electrospray ionization (ESI) source operated in both positive and negative ionization mode. Full-scan mass spectra were acquired over a range of m/z 50–500. ESI parameters were optimized as follows: the spray voltage 3500 V, capillary temperature 350 °C, sheath gas 30 Arb, aux gas 10 Arb, probe heater temp 325 °C. Chromatographic separation was achieved using a C18 column (2.1 mm × 100 mm) with a flow rate of 0.5 mL/min and an injection volume of 5 μL, following the same mobile phase composition and gradient elution program as the HPLC method.

### 4.6 Lipid extraction and quantification

A total of 4 OD units of cell culture were collected into a centrifuge tube and centrifuged to remove the supernatant. To the cell pellet, 100 μL of methyl tridecanoate (C13:0 methyl ester) internal standard was added, followed by 500 μL of 0.5 M sodium-methanol solution. The mixture was vortexed at 1,200 rpm for 2 h. Subsequently, 40 μL of 98% sulfuric acid was added to neutralize the sample. Fatty acid methyl esters (FAMEs) were extracted by adding 400 μL of hexane to each sample, followed by vortexing at 1,200 rpm for 10 min. The samples were then centrifuged at 14,000 rpm for 2 min, and 250 μL of the hexane phase was collected, the hexane phase was filtered through a 0.22 *μ*m membrane filter and analyzed using gas chromatography–flame ionization detection (GC-FID) for quantitative determination of fatty acid content. Total lipid was calculated as the sum of the major fatty acid species detected, including C16:0, C16:1, C17:0, C17:1, C18:0, C18:1, and C18:2.

Statistical analysis of metabolite titer and content was conducted using Student’s t-test, and the significances were defined as: P > 0.05 (ns), P < 0.05 (*), P < 0.01 (**), P < 0.001 (***). Each value represents the mean ± standard deviation from at least three independent experiments (n >3).

## Supporting information

Electronic Supplementary Material

## Author contributions

Jinpeng Wang: Conceptualization, Investigation, Data curation, Methodology, Visualization, Writing - original draft. Yuxiang Hong: Data curation, Visualization, Investigation. Zizhao Wu: Methodology, Investigation. Ayelet Fishman: Writing - review & editing. Peng Xu: Conceptualization, Writing - review & editing, Project administration, Funding acquisition.

## Data availability

Data will be made available on reasonable requests.

## Acknowledgements

This work was financially supported by the National Natural Science Foundation of China (22378083), Guangdong Provincial Key Laboratory of Materials and Technologies for Energy Conversion (MATEC), and Muyuan laboratory (Program ID: 12106022401).

