## Supplementary material for "Engineering amino acid-derived malonyl-CoA pathways to boost polyketide production in *Yarrowia lipolytica*": Electronic Supplementary Material

**Supplementary notes**

**Strain culture conditions.**

**Luria-Bertani (LB) medium:** peptone 10 g/L, sodium chloride 10 g/L, yeast extract 5 g/L, supplemented with 100 mg/mL ampicillin for plasmid construction and extraction.

**Yeast extract peptone dextrose (YPD) medium:** glucose 20 g/L, yeast extract 10 g/L, peptone 20 g/L, for genome extraction or linearized plasmid transformation.

**Fermentation medium (YPD20):** glucose 20 g/L, yeast extract 10 g/L, peptone 20 g/L, for analyzing growth curve of engineered strains.

**Fermentation medium (YPD80):** glucose 80 g/L, yeast extract 10 g/L, peptone 20 g/L, additional supplements (Monopotassium phosphate 4 g/L, Dipotassium phosphate 6 g/L, Calcium chloride 29 mg/L, Magnesium sulfate 0.5 g/L, TRACE elements stock solution 10 mL/L, Ferrous sulfate heptahydrate 0.2 mM, Vitamin stock solution 10 mL/L, Biotin 1 mg/L), for fermentation of engineered strains.

The TRACE elements stock solution contains: EDTA 15 g/L, Zinc sulfate heptahydrate 10.2 g/L, Manganese chloride tetrahydrate 0.5 g/L, Copper sulfate 0.5 g/L, Cobalt chloride hexahydrate 0.86 g/L, Sodium molybdate dihydrate 0.56 g/L.

The vitamin stock solution contains: Calcium pantothenate 1.2 g/L, Nicotinic acid 1.2 g/L, Inositol 30 g/L, p-Aminobenzoic acid 0.24 g/L, Vitamin B1 (Thiamine) 1.2 g/L, Vitamin B6 (Pyridoxine) 1.2 g/L.

**For 1.0 L bioreactor fermentation conditions:**

Initial culture medium: glucose 20 g/L, yeast extract 10 g/L, peptone 20 g/L, additional supplements (Monopotassium phosphate 4 g/L, Dipotassium phosphate 6 g/L, Calcium chloride 29 mg/L, Magnesium sulfate 0.5 g/L, TRACE elements stock solution 10 mL/L, Ferrous sulfate heptahydrate 0.2 mM, Vitamin stock solution 10 mL/L, Biotin 1 mg/L). Antifoam 204 was added to the initial culture medium at a concentration of 0.05% (v/v).

Continuous feed medium: Glucose 500 g/L, YP (a 1:2 mixture of yeast extract and peptone) 300 g/L. The pH of the fermentation broth was maintained at 6 using HCl (4 M) and NaOH (4 M).

**Synthetic drop-out medium without uracil (SD-Ura):** glucose 20 g/L, amino acid-free and ammonium sulfate-free YNB 1.7 g/L, ammonium sulfate 5 g/L, complete supplement mixture-Ura (CSM-Ura): 0.67 g/L, for isolating positive *Y. lipolytica* transformants.

**Synthetic drop-out medium without uracil (SD-Leu):** glucose 20 g/L, amino acid-free and ammonium sulfate-free YNB 1.7 g/L, ammonium sulfate 5 g/L, complete supplement mixture-Leu (CSM-Leu): 0.67 g/L, for isolating positive *Y. lipolytica* transformants.

**YPD medium containing 5-fluoroorotic acid (YPD-5-FOA):** Add 2.5 mL of 5-fluoroorotic acid at a concentration of 100 mg/mL to 250 mL of YPD medium, used to eliminate the URA3 selection marker.

**Supplementary Figures**

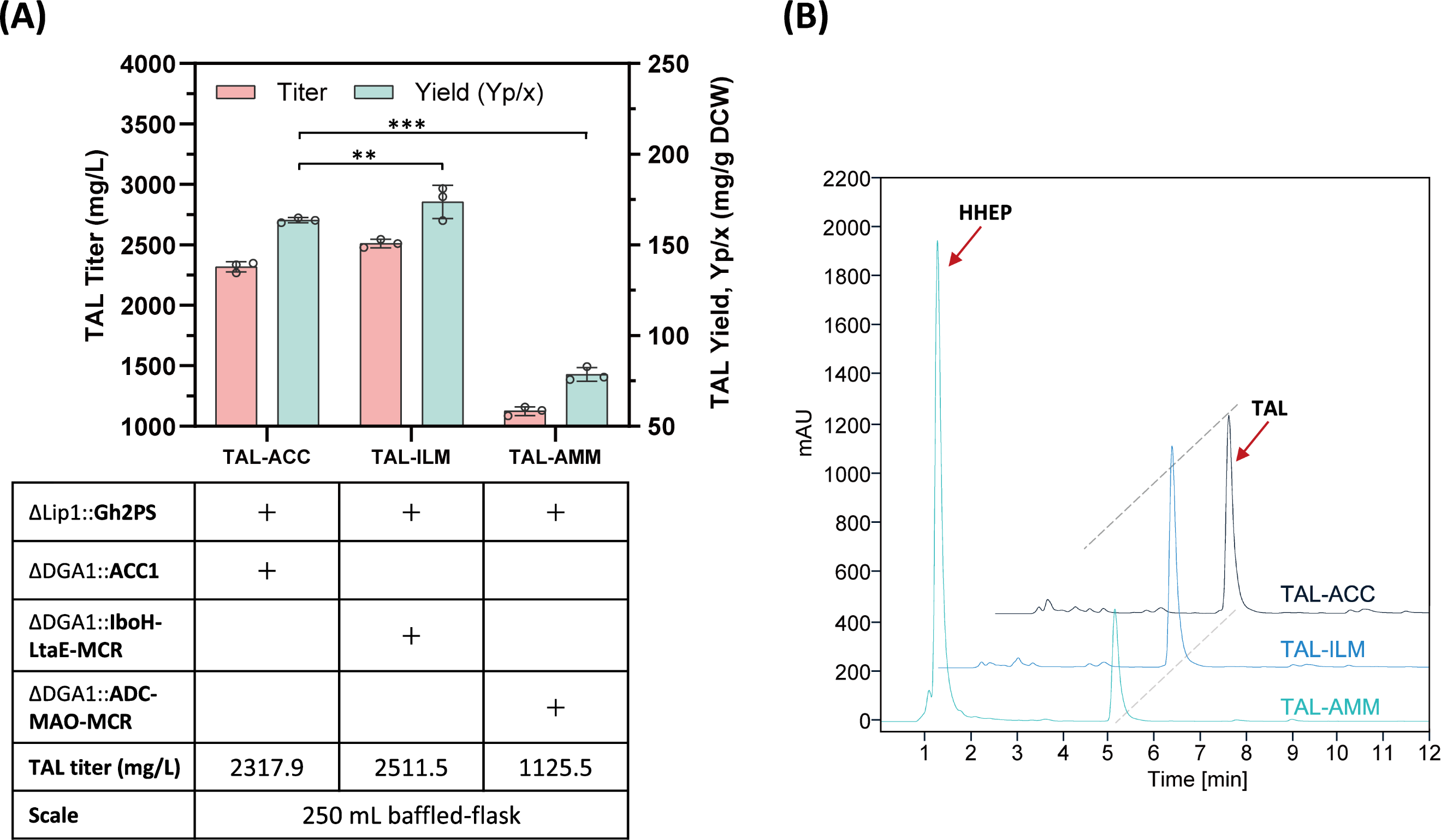

**Supplementary Fig. S1**. Testing the functionality of the novel malonyl-CoA pathway in shaking flasks with baffles. (**A**) TAL production from the novel malonyl-CoA pathways. (**B**) HPLC profile persists of the unknown peak when AMM pathway was employed.

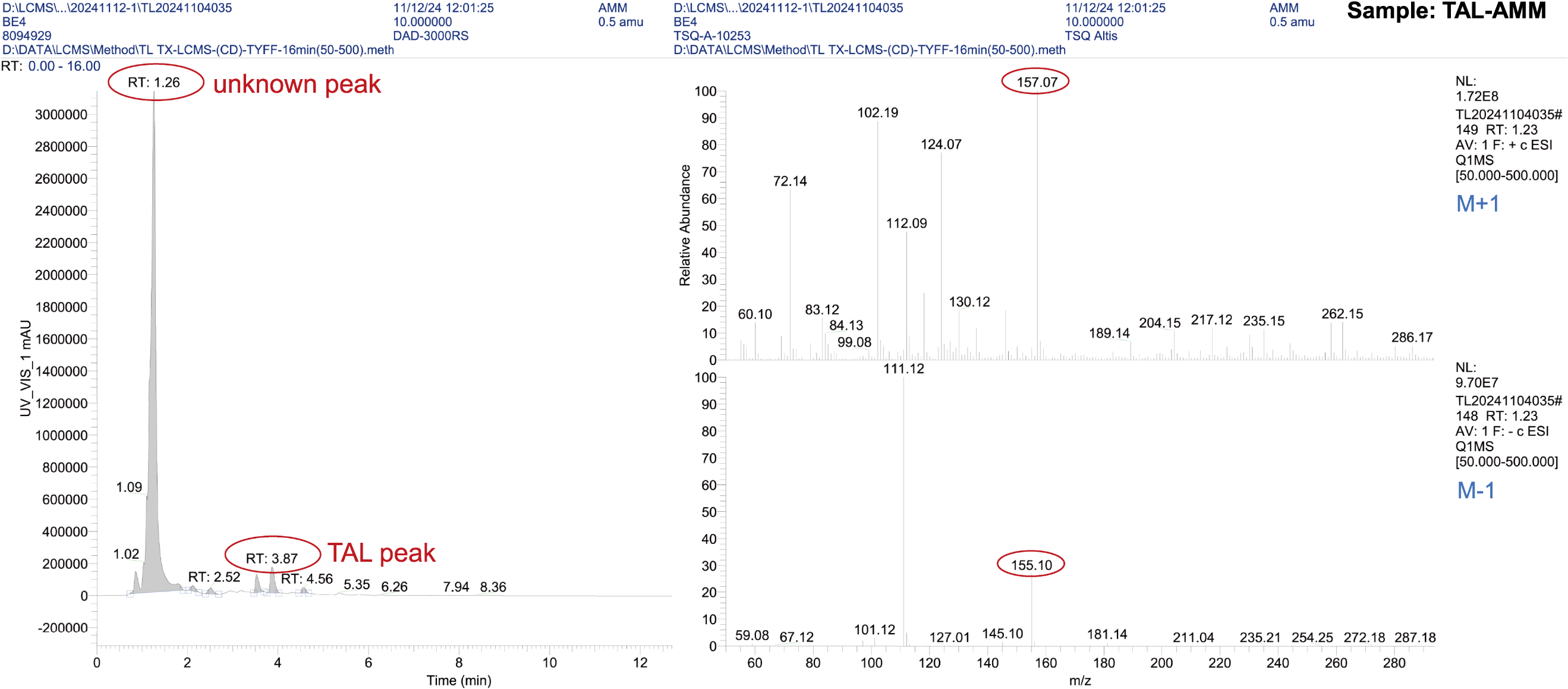

**Supplementary Fig. S2**. LC-MS profile of the unknown peak at 1.26 min and the mass fragmentation pattern of the unknown compound. 3.87 min is the TAL peak.

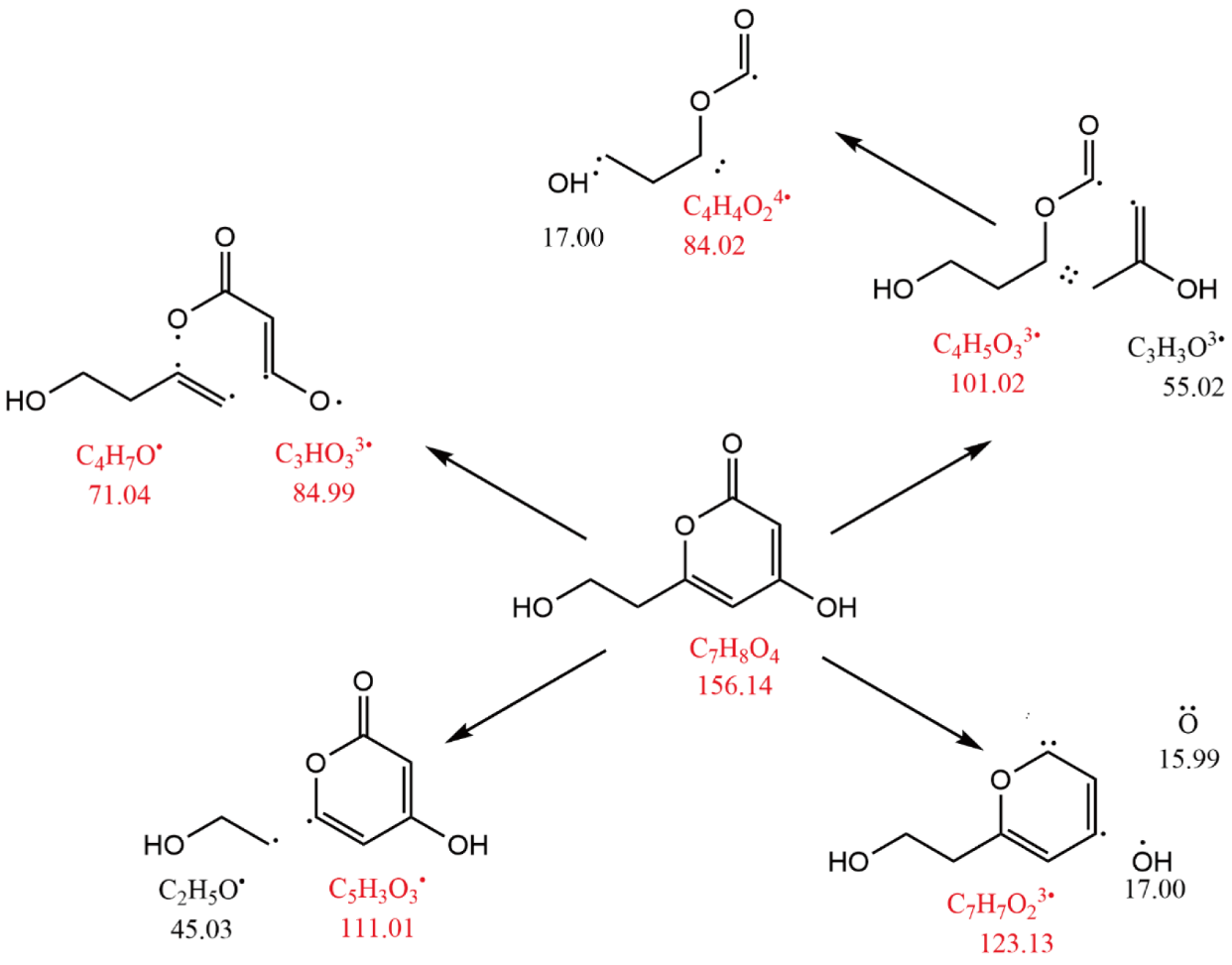

**Supplementary Fig. S3**. Matching the mass fragmentation pattern of the unknown peak at m/z of 156, 123, 111, 101, 84 and 71.

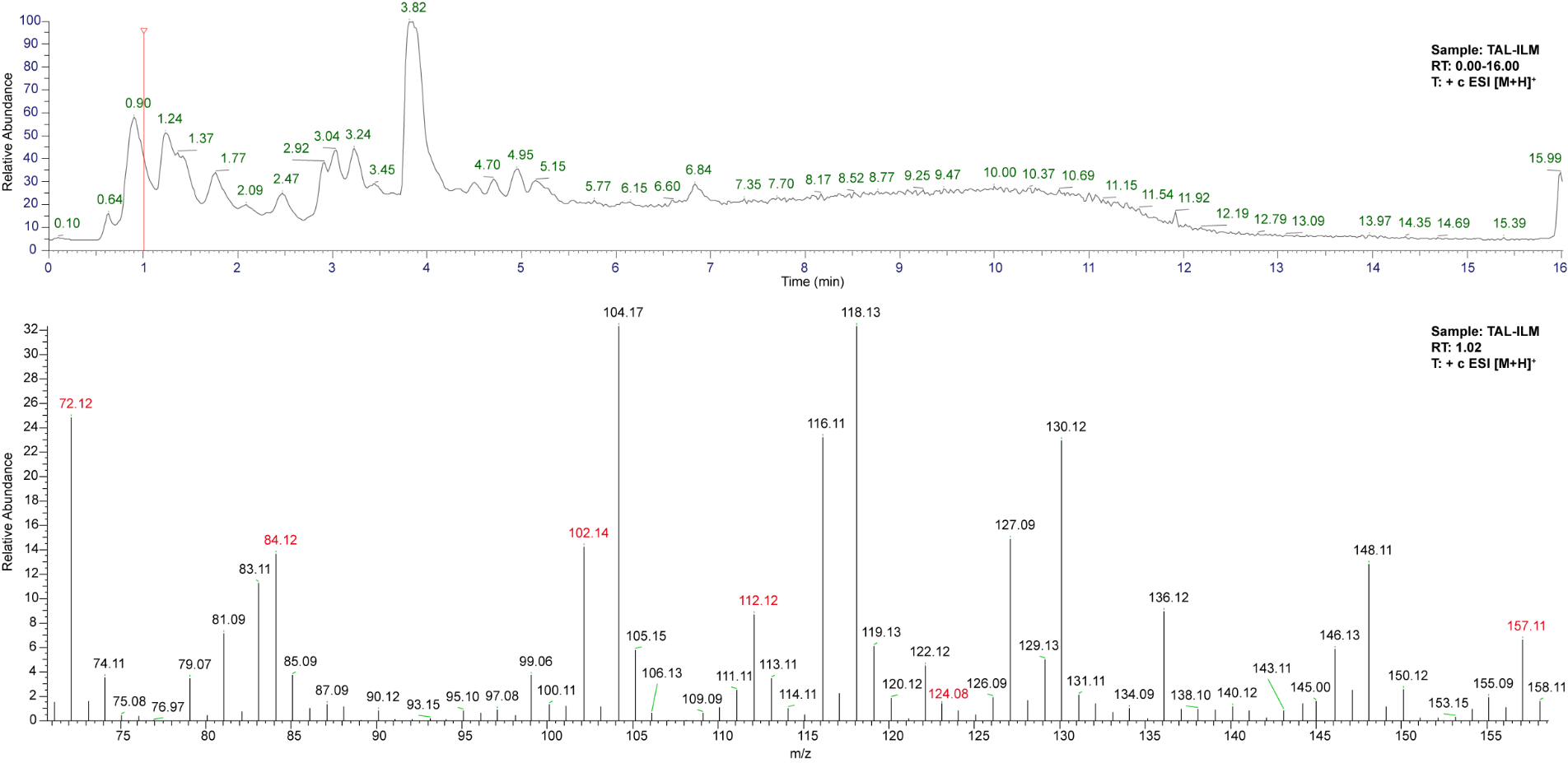

**Supplementary Fig. S4**. LC-MS profile of the unknown peak at around 1.02 min in the TAL-ILM strain (upper panel) and the appearance of m/z 157, 124, 112, 102, 84 and 72 of the unknown compound (lower panel). The m/z of the unknown peak is relatively diminished by the noise peak (lower panel). 3.82 min is the TAL peak.

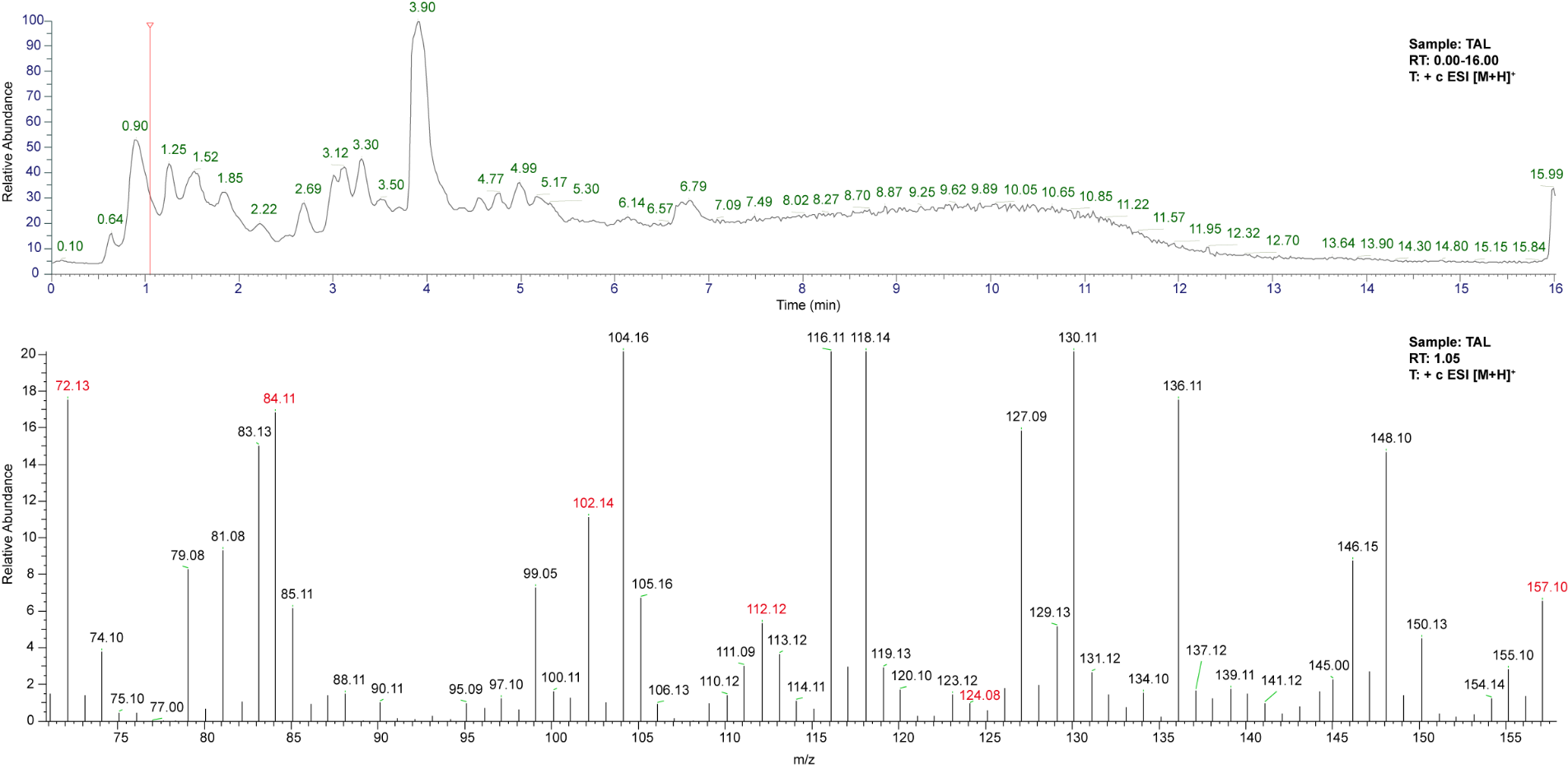

**Supplementary Fig. S5**. LC-MS profile of the unknown peak at around 1.05 min in the TAL strain (upper panel) and the appearance of m/z 157, 124, 112, 102, 84 and 72 of the unknown compound (lower panel). The m/z of the unknown peak is relatively diminished by the noise peak (lower panel). 3.90 min is the TAL peak.

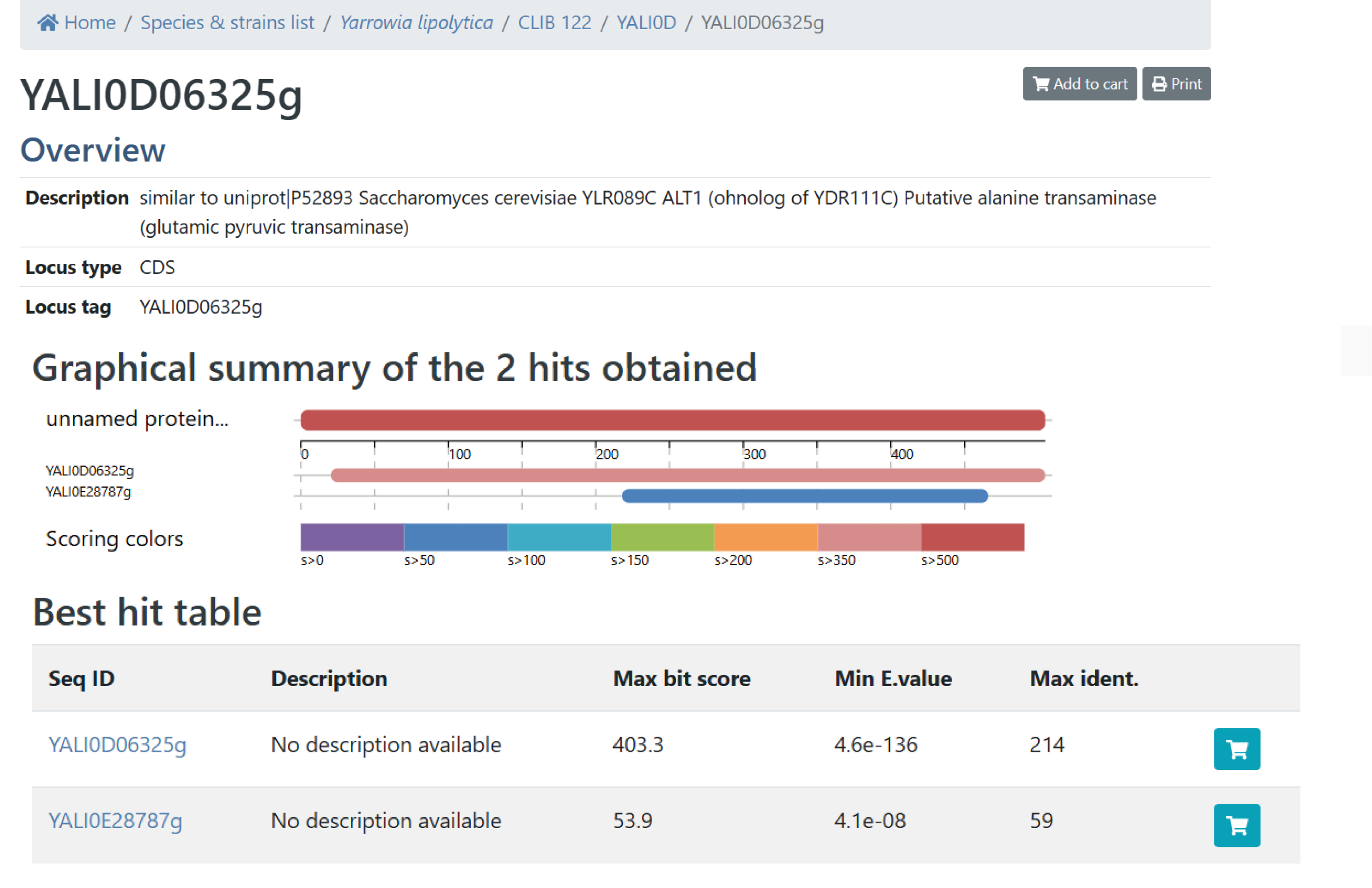

**Supplementary Fig. S6**. Bioinformatic blasting confirms the presence of YAlI0D06325 as the putative β-alanine-pyruvate aminotransferase (BAPAT) in *Y. lipolytica* genome. The blasting input protein is human alanine aminotransferase 1 annotated as GPT with a Uniprot ID P24298. The blasting was done using the GRYC database (https://gryc.inrae.fr).

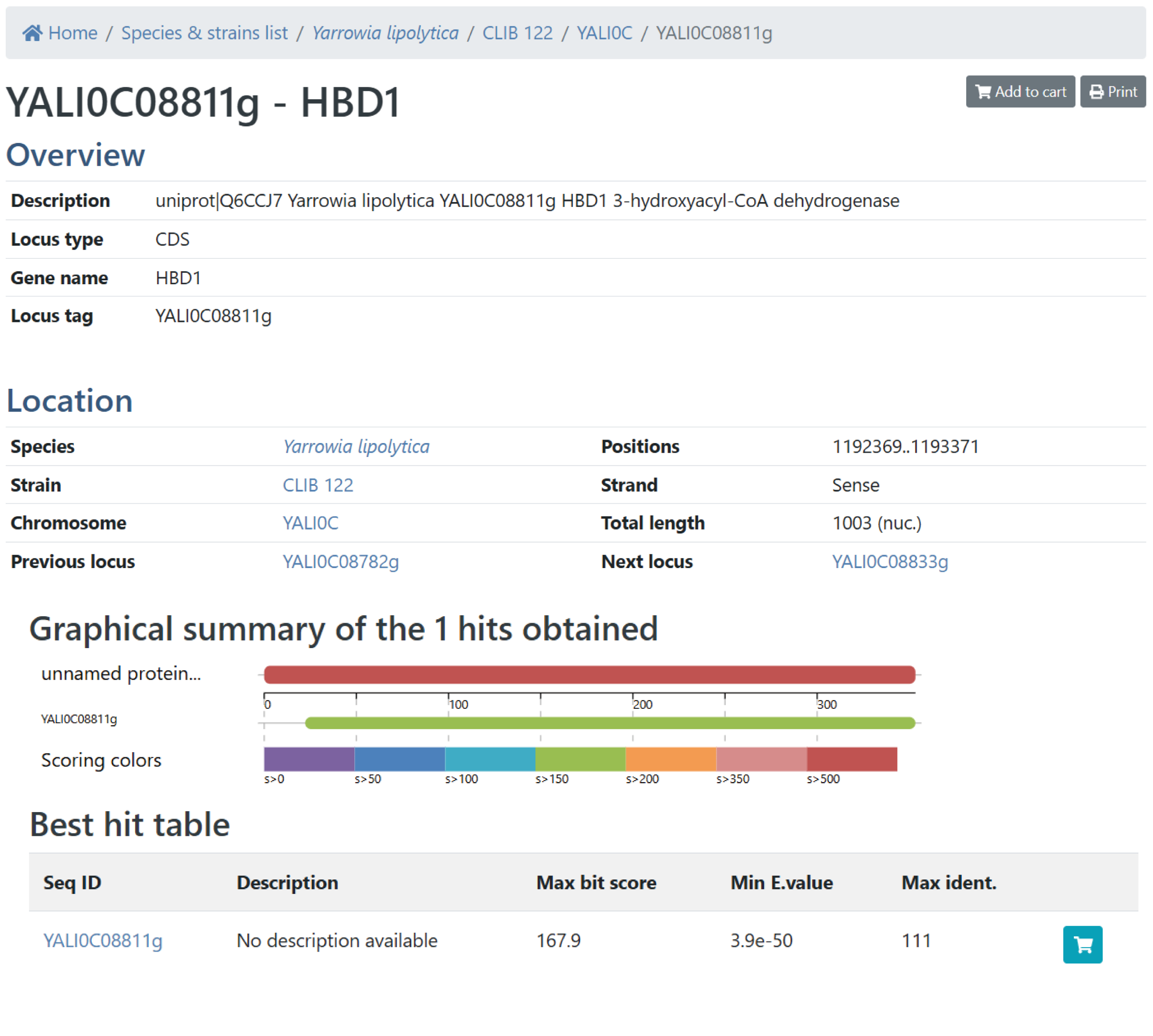

**Supplementary Fig. S7**. Bioinformatic blasting confirms the presence of YALI0C08811 as the putative 3-hydroxyacyl-CoA dehydrogenase / 3-hydroxypropionate dehydrogenase (HBD1) in *Y. lipolytica* genome. The blasting input protein is annotated as HBD1 with a uniport ID B3KTT6. The blasting was done using the GRYC database (https://gryc.inrae.fr).

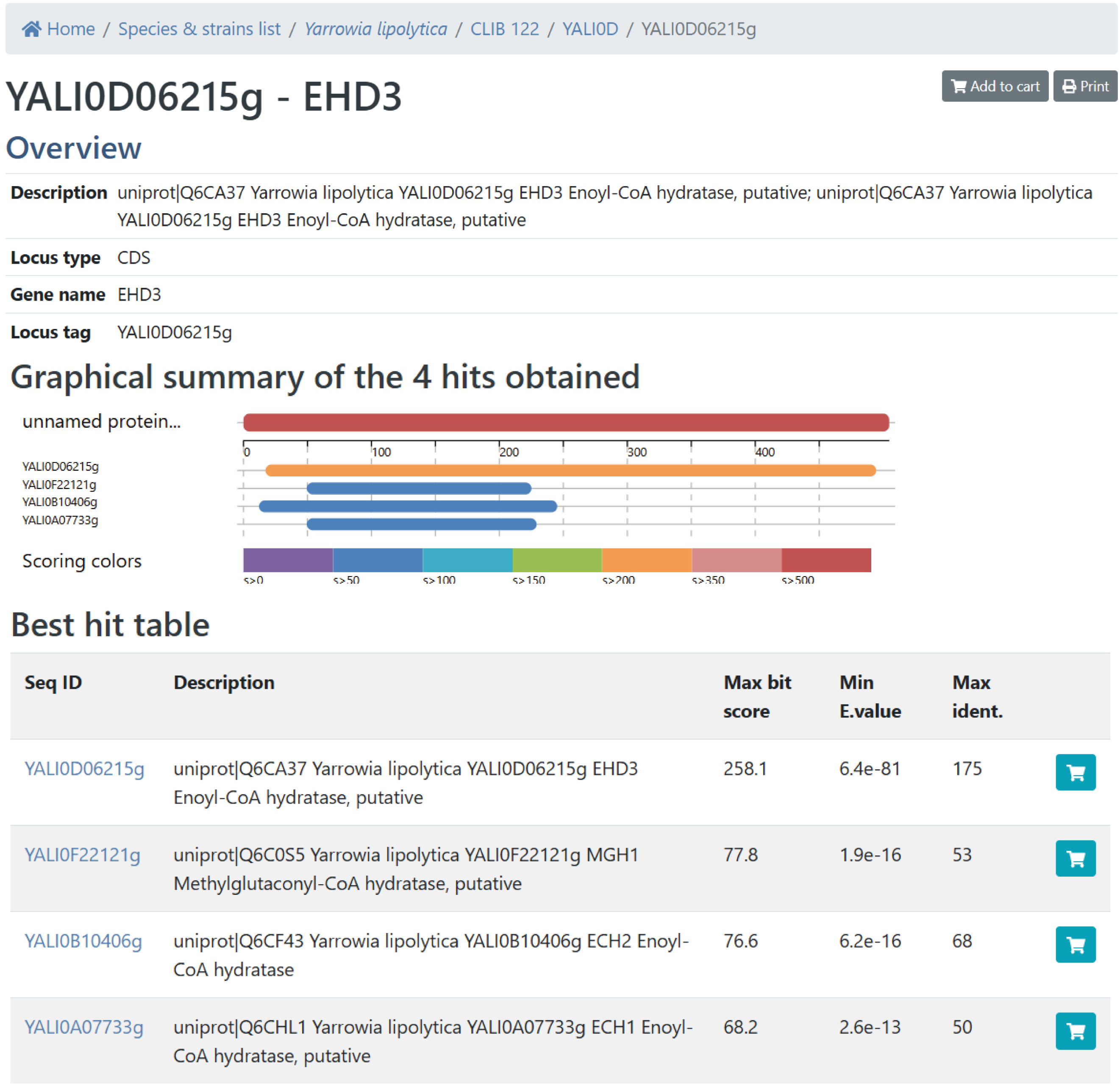

**Supplementary Fig. S8**. Bioinformatic blasting confirms the presence of YALI0D06215 as the putative enoyl-CoA hydratase/3-hydroxyisobutyryl-CoA hydrolase (EHD3) in *Y. lipolytica* genome. The blasting input protein is annotated as EHD3 with a Uniprot ID P28817. The blasting was done using the GRYC database (https://gryc.inrae.fr).

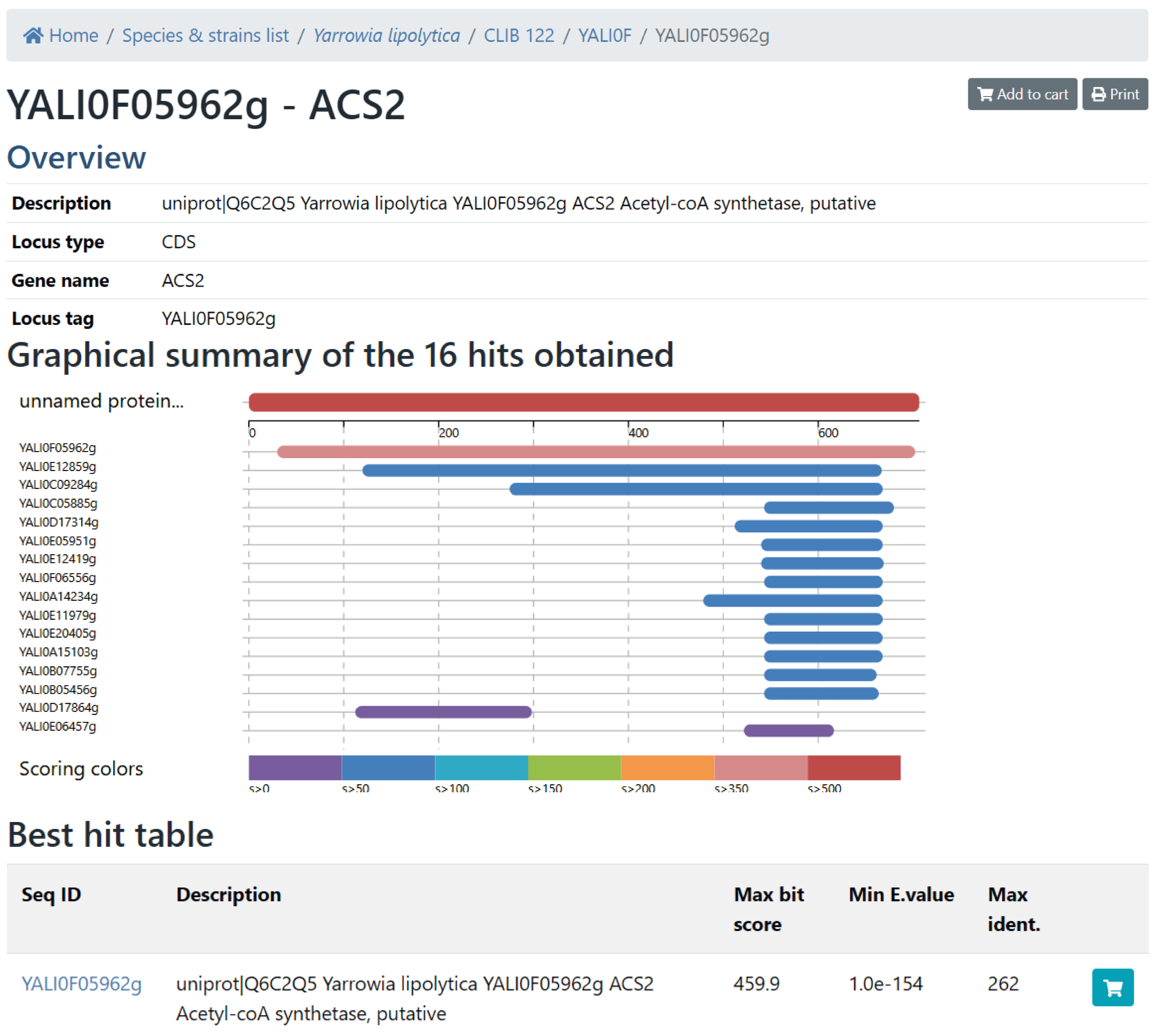

**Supplementary Fig. S9**. Bioinformatic blasting confirms the presence of YALI0F05962 as the putative acetyl-CoA synthetase/3-hydroxypropionyl-CoA synthetase (ACS2) in *Y. lipolytica* genome. The blasting input protein is annotated as 3-hydroxypropionyl-CoA synthetase with a Uniprot ID A4YGR1. The blasting was done using the GRYC database (https://gryc.inrae.fr).

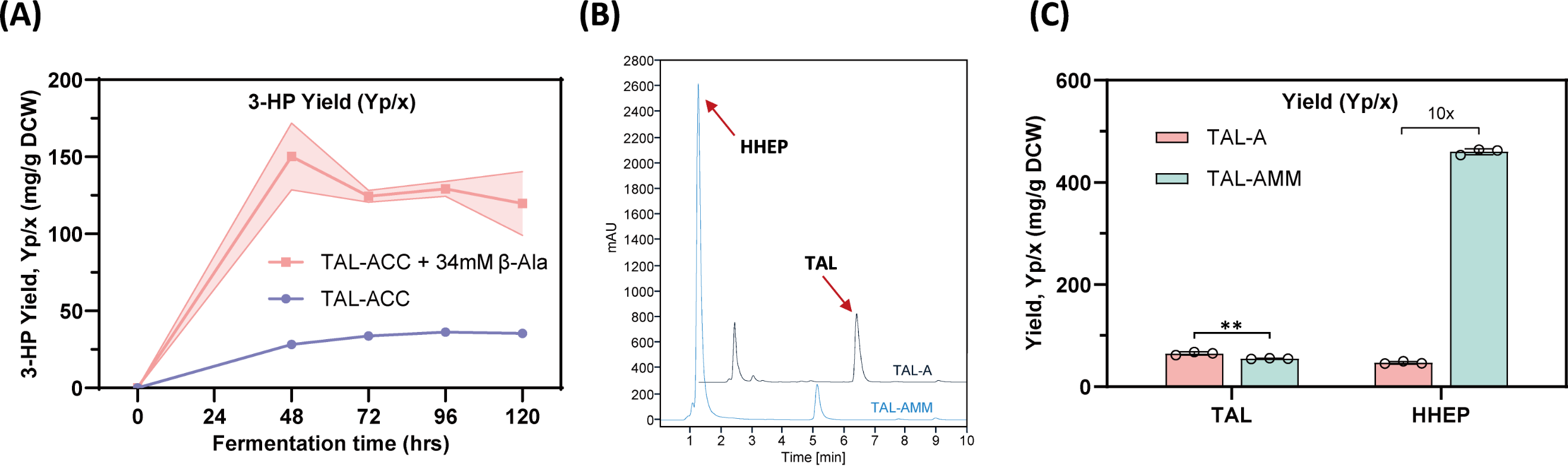

**Supplementary Fig. S10**. Identification of the synthetic pathway for HHEP with feeding of β-alanine to the TAL-ACC strain. (**A**) 3-HP production after feeding β-alanine into TAL-ACC strain. (**B**) HPLC profile persists of the HHEP peak when *Sf*ADC was expressed. (**C**) Comparison of TAL and HHEP yield when *Sf*ADC or entire AMM pathway was expressed.

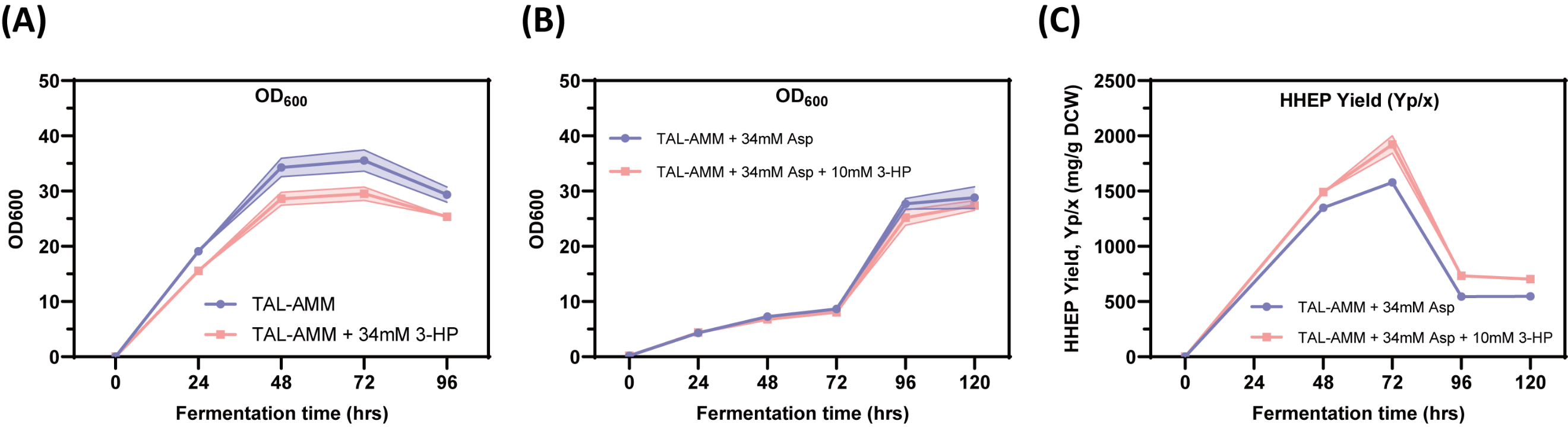

**Supplementary Fig. S11**. Identification of the synthetic pathway for HHEP with feeding of 3-HP to the TAL-AMM strain. (**A**) Strain growth curves with or without feeding 34 mM 3-HP. 3-HP feeding: 5 mM, 5 mM, 8 mM, 8 mM, and 8 mM were added at 0 h, 12 h, 24 h, 36 h, and 48 h, respectively, resulting in a total supplement of 34 mM. (**B**) Strain growth curves with or without feeding 10 mM 3-HP. 3-HP feeding: 1.4 mM, 1.4 mM, 2.4 mM, 2.4 mM, and 2.4 mM were added at 24 h, 36 h, 48 h, 60 h, and 72 h, respectively, resulting in a total supplement of 10 mM. L-Aspartate feeding: 5 mM, 5 mM, 8 mM, 8 mM, and 8 mM were added at 24 h, 36 h, 48 h, 60 h, and 72 h, respectively, resulting in a total supplement of 34 mM. (**C**) HHEP yield on biomass.

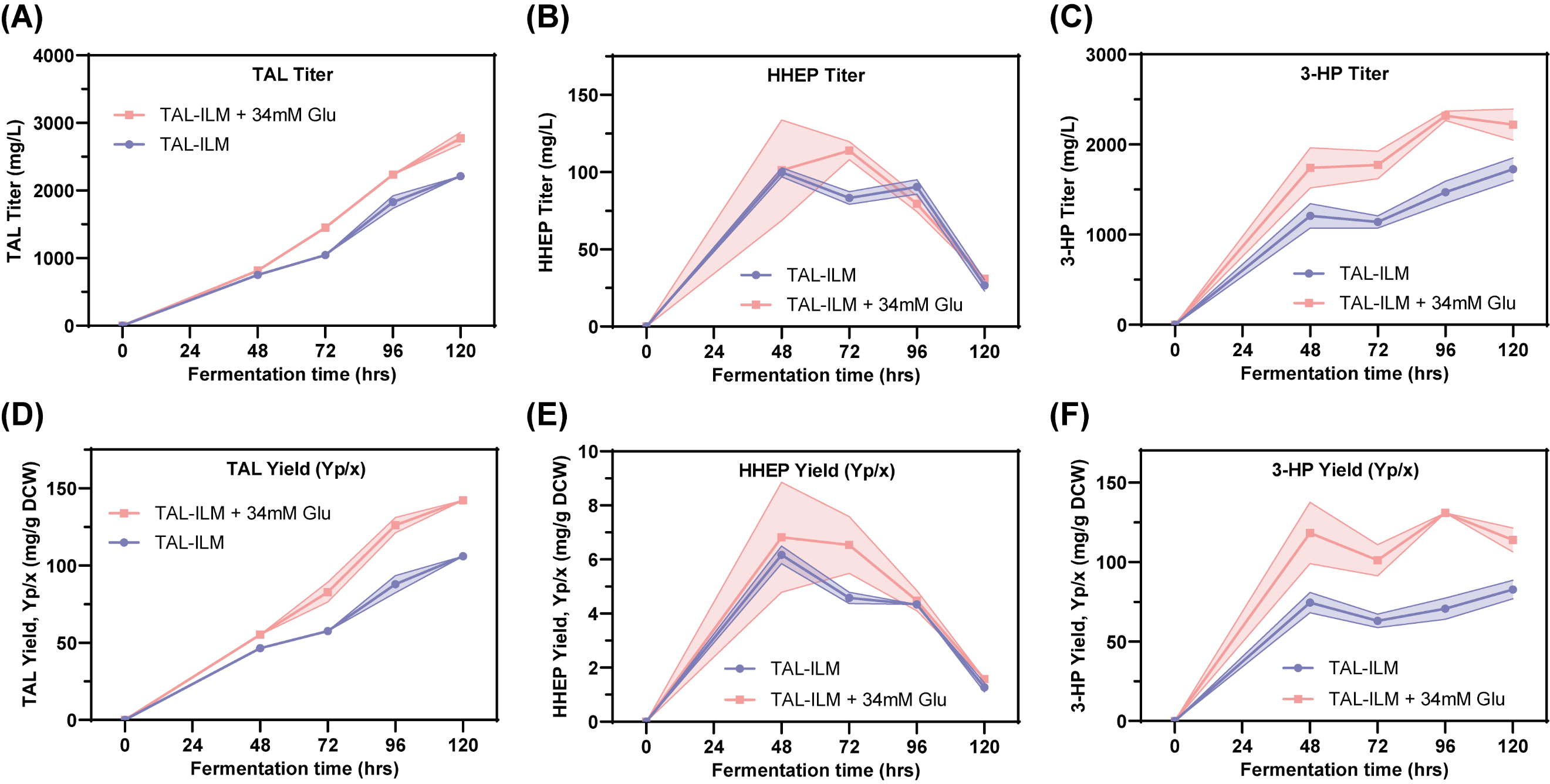

**Supplementary Fig. S12**. Testing the functionality of the TAL-ILM pathway by feeding 34 mM glutamate in shaking flask. (**A**) TAL titer. (**B**) HHEP titer. (**C**) 3-HP titer. (**D**) TAL yield based on biomass. (**E**) HHEP yield on biomass. (**F**) 3-HP yield on biomass.

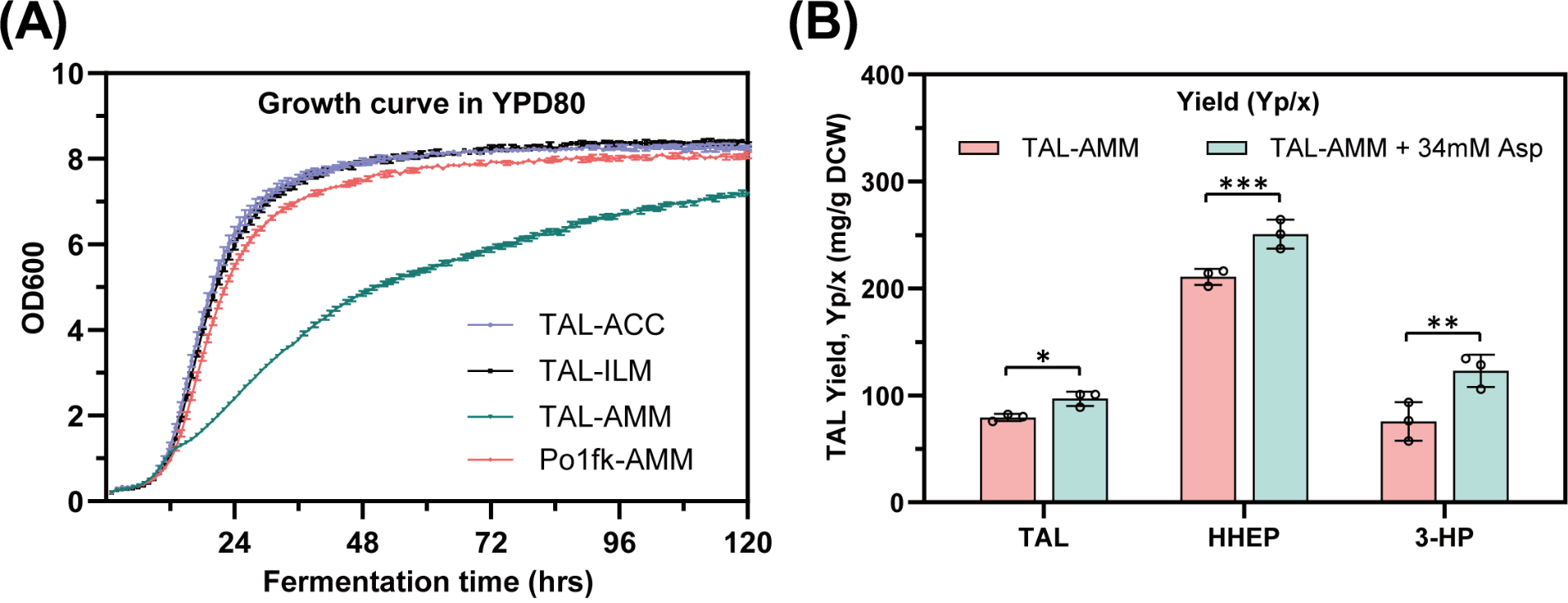

**Supplementary Fig. S13**. (**A**) The growth curve of the engineered strains was monitored in YPD80 medium by an automatic growth analyzer. (**B**) Testing the functionality of the TAL-AMM pathway by feeding 34 mM aspartate in shaking flask, the yield of TAL, HHEP and 3-HP was calculated on the basis of biomass.

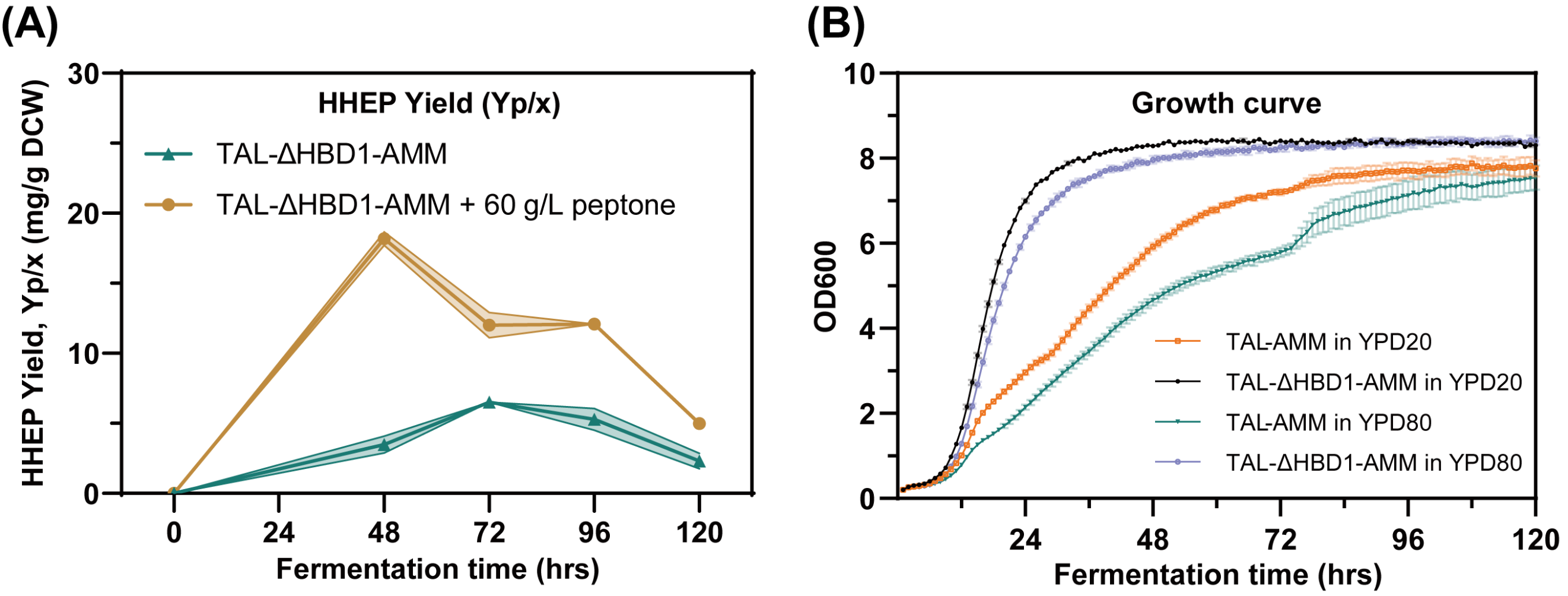

**Supplementary Fig. S14**. (**A**) Production of HHEP in the TAL-ΔHBD1-AMM strain with or without feeding of 60 g/L peptone in shaking flask using YPD80 medium. (**B**) The growth curve of the engineered strains was monitored in YPD20 or YPD80 medium by an automatic growth analyzer.

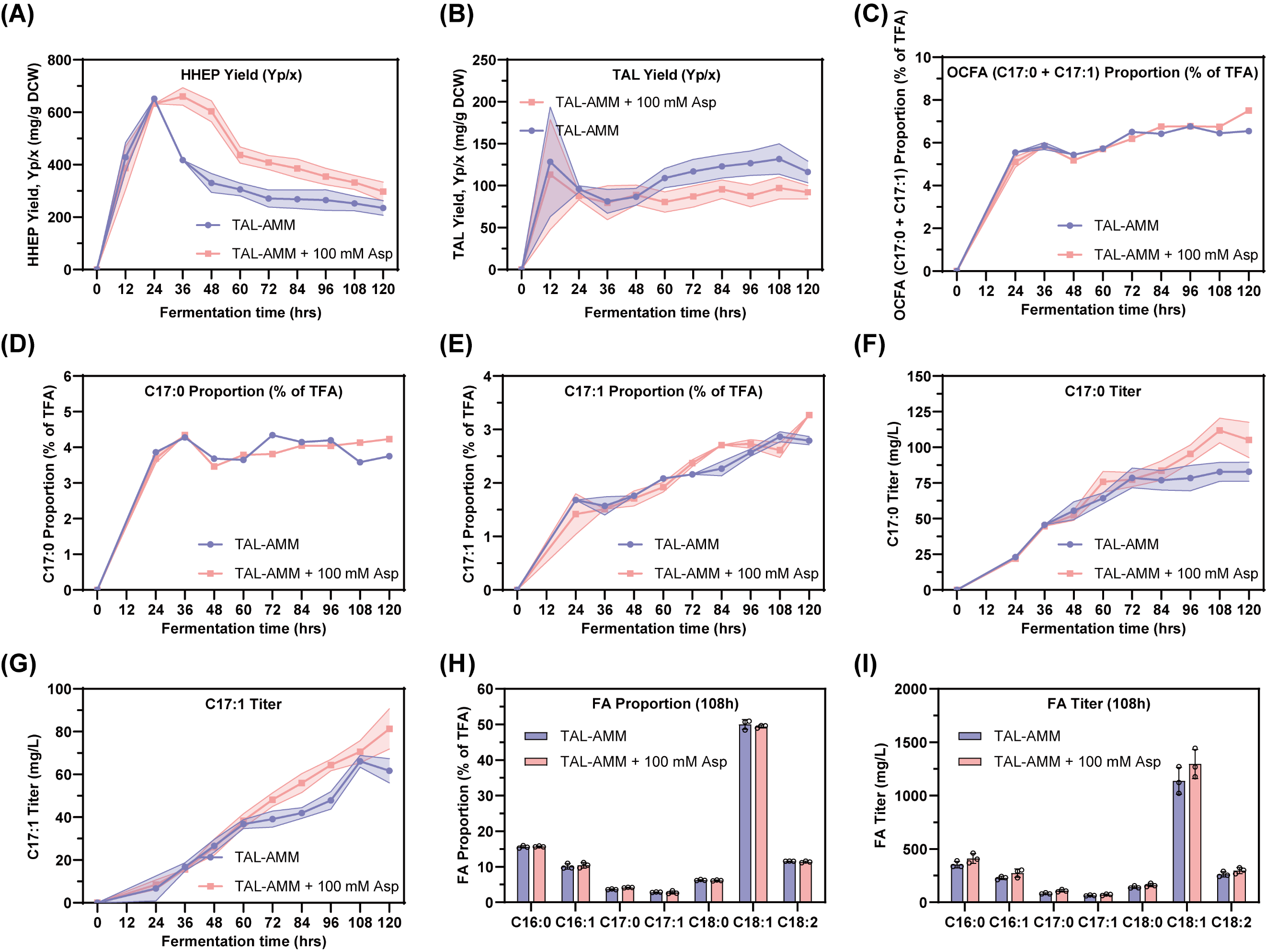

**Supplementary Fig. S15**. Feeding L-aspartate to improve polyketide and lipid production in TAL-AMM strain in a 1.0 L mini-reactor. **(A)** HHEP yield on biomass during fermentation; **(B)** TAL yield on biomass during fermentation; **(C)** The proportion of total odd-chain fatty acids (C17:0 and C17:1) during fermentation; **(D)** The proportion of heptadecanoic acid (C17:0) during fermentation; **(E)** The proportion of heptadecenoic acid (C17:1) during fermentation; **(F)** The titer of heptadecanoic acid (C17:0) during fermentation; **(G)** The titer of heptadecenoic acid (C17:1) during fermentation; **(H)** The proportion of total fatty acids in 108h; **(I)** The titer of total fatty acids in 108h.

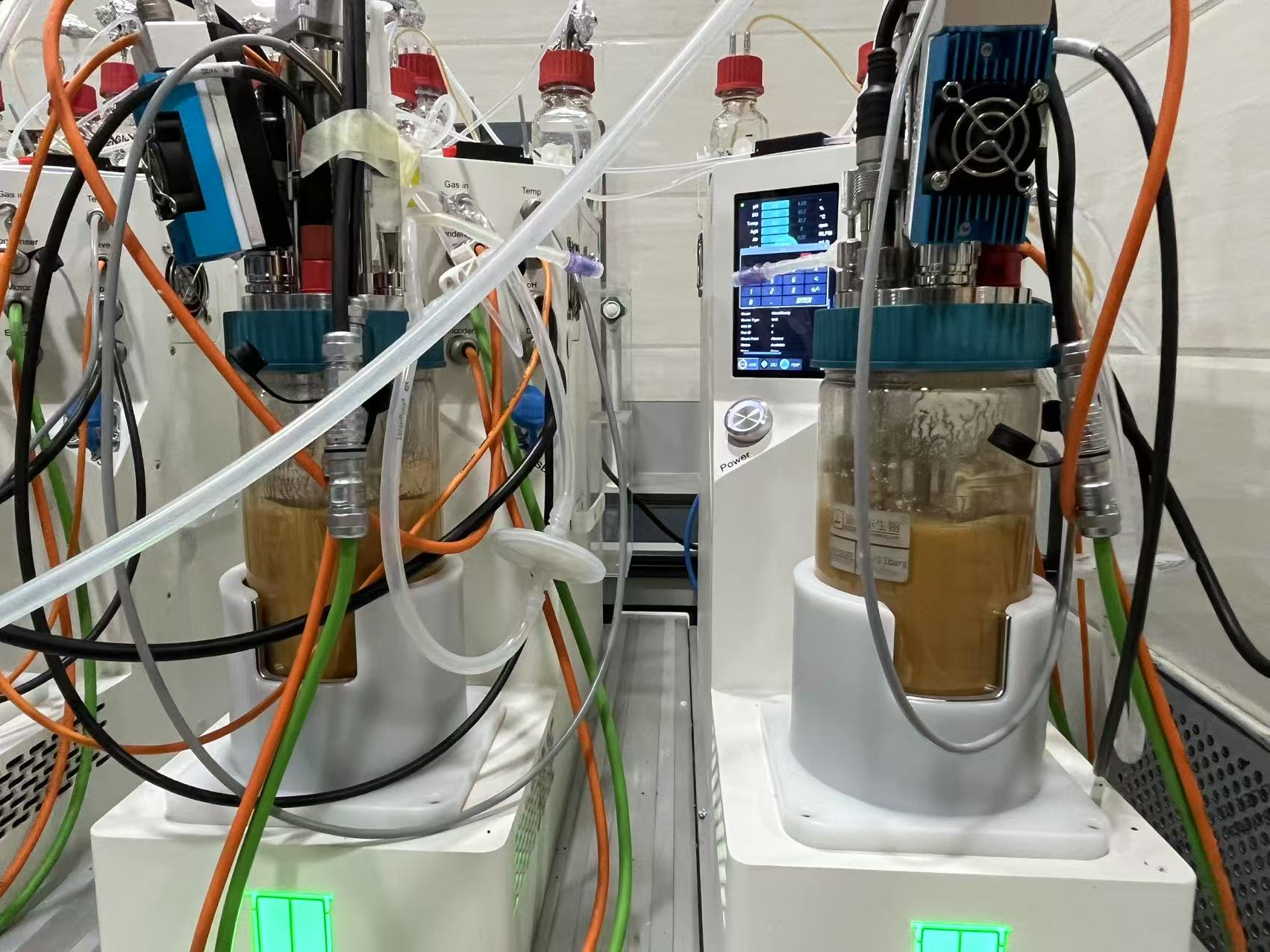

**Supplementary Fig. S16**. Feeding L-aspartate to improve polyketide and lipid production in TAL-AMM strain in a 1.0 L mini-reactor. The left bioreactor was fed with 100 mM L-aspartate. The right bioreactor was not fed.

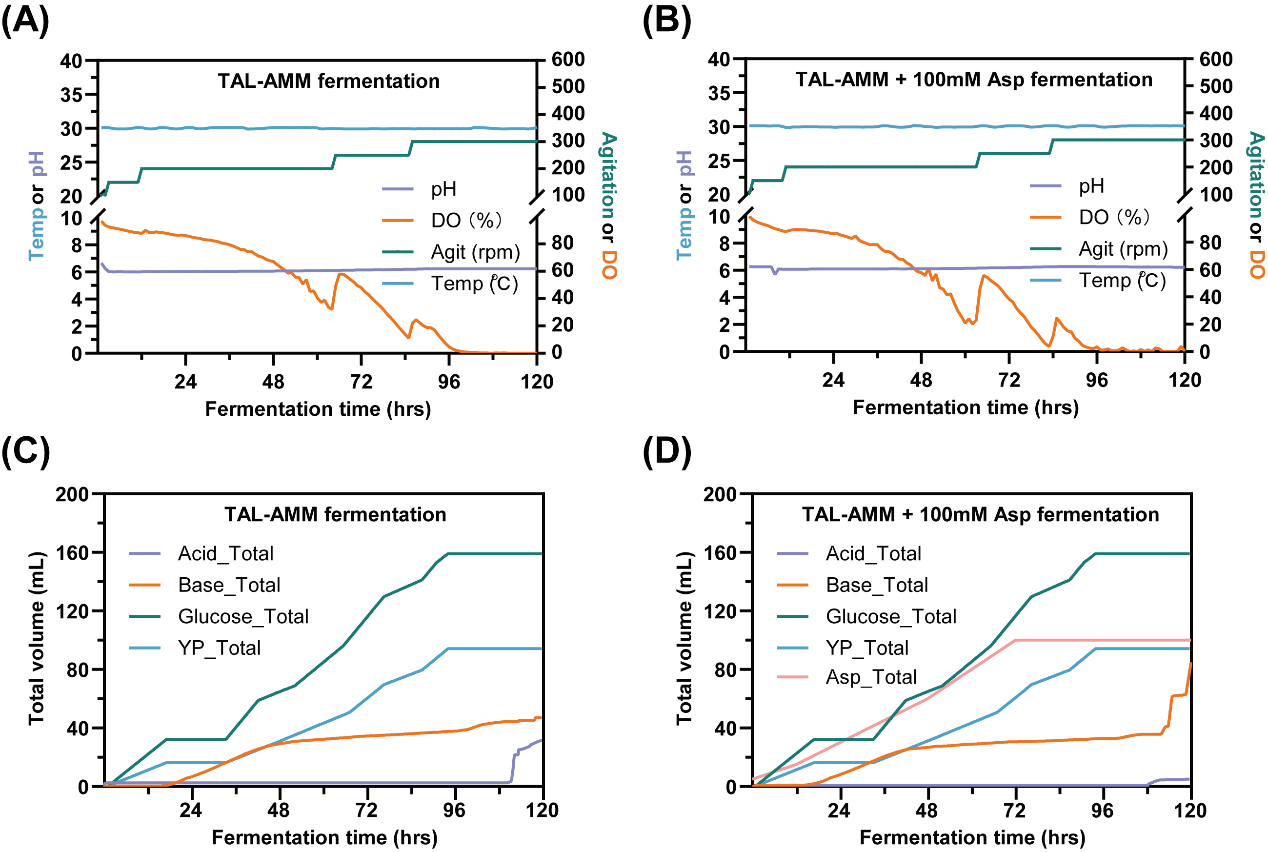

**Supplementary Fig. S17**. Feeding L-aspartate to improve polyketide and lipid production in TAL-AMM strain in a 1.0 L mini-reactor.

(A) and (B) Bioreactor fermentation process parameters.

(C) Feeding profile for the non-fed group during bioreactor operation: HCl (4 M), total feed 32 mL; NaOH (4 M), total feed 47 mL; Glucose (500 g/L), total feed 160 mL; YP (a 1:2 mixture of yeast extract and peptone, 300 g/L), total feed 95 mL.

(D) Feeding profile for the fed group during bioreactor operation: HCl (4 M), total feed 5 mL; NaOH (4 M), total feed 85 mL; Glucose (500 g/L), total feed 160 mL; YP (a 1:2 mixture of yeast extract and peptone, 300 g/L), total feed 95 mL; L-Aspartate feeding: 5 mM, 10 mM, 15 mM, 15 mM, 15 mM, 20 mM, and 20 mM were added at 0 h, 12 h, 24 h, 36 h, 48 h, 60 h, and 72 h, respectively, resulting in a total supplement of 100 mM.

**Supplementary Tables**

**Table S1. Plasmids used or constructed in this study.**

| Plasmids | Genotype or relevant properties | Source |
| --- | --- | --- |
| pYLXP' | Amp+, *E. coli* and *Y. lipolytica* shuttle vector, YaliBrick plasmid, Leu+, pTEFin promoter, XPR2 terminator | [1] |
| pUrLp | Amp+, based on pYLXP', the pTEFin promoter and XPR2 terminator were replaced by loxP-Ura3-loxP cassette | [2] |
| pUC-Leu-A08 | Amp+, Leu^+^, A08 5'-upstream and 3'-downstream homology arms inserted into pUC57, used for integrating gene in the A08 (YALI0A08382) locus in *Y. lipolytica* | [3] |
| pYL31 | Amp+, Replacing the P_TEFin_ promoter and T_Xpr2_ terminator region of pYLXP' with P_FBA_ promoter and T_Pex20_ terminator, Leu^+^ | This study |
| pYL24 | Amp+, Replacing the P_TEFin_ promoter and T_Xpr2_ terminator region of pYLXP' with P_TDH1_ promoter and T_Mig1_ terminator, Leu^+^ | This study |
| pYLXP'-*Gh*2PS | Amp+, P_TEFin_-*Gh2PS*-xpr2t, used for subcloning | This study |
| pYLXP'-*Yl*ACC1 | Amp+, P_TEFin_-*YlACC1*-xpr2t, used for subcloning | This study |
| pYL31-*Ps*LtaE | Amp+, P_FBA_-*PsLtaE*-pex20t, used for subcloning | This study |
| pYL24-*Am*IboH | Amp+, P_TDH1_-*AmIboH*-mig1t, used for subcloning | This study |
| pYLXP'-*Ca*MCR-C | Amp+, P_TEFin_-*CaMCR-C*-xpr2t, used for subcloning | This study |
| pYL31-*Sf*ADC | Amp+, P_FBA_-*SfADC*-pex20t, used for subcloning | This study |
| pYL24-*Ec*MAO | Amp+, P_TDH1_-*EcMAO*-mig1t, used for subcloning | This study |
| pUrLp-ΔLip1 | Amp+, Lip1 5'-upstream and 3'-downstream homology arms inserted into pUrLp, used for integrating gene in the Lip1 (YALI0E10659g) locus in *Y. lipolytica* | This study |
| pUrLp-ΔDga1 | Amp+, Dga1 5'-upstream and 3'-downstream homology arms inserted into pUrLp, used for integrating gene in the Dga1 (YALI0E32769g) locus in *Y. lipolytica* | This study |
| pUrLp-ΔHBD1 | Amp+, HBD1 5'-upstream and 3'-downstream homology arms inserted into pUrLp, used for deleting HBD1 gene (YALI0C08811g) in the in *Y. lipolytica* | This study |
| pUrLp-ΔLip1-*Gh*2PS | Amp+, loxP-Ura3-loxP cassette, Lip1-up and Lip1-down, P_TEFin_-*Gh2PS*-xpr2t, used for integrating *Gh2PS* in the Lip1 locus in *Y. lipolytica* | This study |
| pUrLp-ΔDga1-*Yl*ACC1 | Amp+, loxP-Ura3-loxP cassette, Dga1-up and Dga1-down, P_TEFin_-*YlACC1*-xpr2t, used for integrating *YlACC1* in the Dga1 locus in *Y. lipolytica* | This study |
| pUrLp-ΔDga1-*Am*IboH-*Ps*LtaE-*Ca*MCR-C | Amp+, loxP-Ura3-loxP cassette, Dga1-up and Dga1-down, P_TDH1_-*AmIboH*-mig1t, P_FBA_-*PsLtaE*-pex20t, P_TEFin_-*CaMCR-C*-xpr2t, used for integrating *AmIboH*, *PsLtaE* and *CaMCR-C* in the Dga1 locus in *Y. lipolytica* | This study |
| pUrLp-ΔDga1-*Sf*ADC | Amp+, loxP-Ura3-loxP cassette, Dga1-up and Dga1-down, P_FBA_-*SfADC*-pex20t, used for integrating *SfADC* in the Dga1 locus in *Y. lipolytica* | This study |
| pUrLp-ΔDga1-*Sf*ADC-*Ec*MAO-*Ca*MCR-C | Amp+, loxP-Ura3-loxP cassette, Dga1-up and Dga1-down, P_FBA_-*SfADC*-pex20t, P_TDH1_-*EcMAO*-mig1t, P_TEFin_-*CaMCR-C*-xpr2t, used for integrating *SfADC*, *EcMAO* and *CaMCR-C* in the Dga1 locus in *Y. lipolytica* | This study |

**Table S2. Synthetic gene sequences used in this study.**

| Names | Sequences (5’ — 3’) |
| --- | --- |
| Codon optimized 2-pyrone synthase from *Gerbera hybrida* coding DNA (*Gh2PS*) for *Y. lipolytica* based on UniProt ID: P48391 | ATGGGGTCATATAGCAGTGATGATGTGGAGGTGATCCGCGAGGCTGGCCGCGCACAAGGACTGGCTACTATCCTGGCCATTGGAACCGCCACACCACCAAATTGCGTGGCTCAAGCCGATTACGCAGACTATTATTTCCGGGTAACTAAATCAGAGCACATGGTCGATCTGAAGGAGAAGTTTAAAAGGATTTGCGAAAAGACTGCCATTAAAAAGCGATATCTCGCCCTCACGGAAGATTATCTCCAGGAAAACCCAACCATGTGTGAGTTCATGGCTCCCTCATTGAACGCTCGCCAGGATCTCGTGGTTACCGGGGTCCCCATGCTTGGCAAGGAGGCCGCCGTCAAAGCCATCGACGAATGGGGTTTGCCAAAGTCAAAAATTACCCATCTGATTTTCTGCACTACCGCCGGGGTAGACATGCCCGGTGCAGACTACCAGCTGGTGAAGCTGCTGGGTCTTTCCCCATCTGTGAAGCGCTATATGCTGTACCAGCAGGGCTGTGCAGCTGGTGGTACTGTGCTGCGCCTGGCTAAGGACTTGGCAGAGAACAATAAGGGGTCACGGGTGCTGATCGTCTGCTCCGAGATTACAGCCATCCTGTTTCACGGACCAAACGAGAATCACCTCGACTCACTGGTGGCTCAAGCTCTGTTCGGCGACGGTGCTGCCGCGCTGATAGTGGGGTCAGGGCCCCATCTGGCTGTGGAACGGCCCATCTTTGAGATTGTTAGCACAGATCAGACCATCTTGCCCGACACTGAGAAAGCGATGAAGCTTCACTTGAGGGAGGGGGGTCTCACGTTCCAGCTCCATAGGGACGTGCCACTGATGGTTGCAAAAAACATCGAAAACGCTGCCGAGAAAGCGCTGTCTCCATTGGGGATTACAGACTGGAACTCTGTGTTTTGGATGGTTCACCCTGGAGGGAGAGCAATCCTGGATCAGGTGGAGCGCAAACTGAACCTTAAAGAGGACAAACTGAGAGCCAGCAGACACGTGCTGAGCGAGTATGGAAACTTGATTTCTGCTTGCGTGCTTTTTATCATCGACGAGGTGCGCAAGCGCTCCATGGCCGAAGGTAAGAGCACTACCGGGGAGGGGCTGGATTGTGGAGTGCTCTTTGGATTTGGTCCAGGTATGACCGTCGAAACTGTTGTACTCCGATCCGTTCGAGTGACCGCTGCAGTGGCCAACGGCAACTAA  (1209 bp) |
| Codon optimized 2-oxoglutarate-dependent dioxygenase from *Amanita muscaria* coding DNA (*AmIboH*) for *Y. lipolytica* based on UniProt ID: A0A0C2SRU5 | ATGCCCCCCACCATCCGAAACAACTTCCCCCTGTTCCACGACTTCGTGACCGCCTCTCACTTCATCACTCAGACCATCCTGATGCGACTGTCTGACGCCATGTACCTGGAGGGCTCTACCCGATTCGAGAACTCTCACCGAGAGGACAAGCCCTCTACCACTACCCTGGTGCTGCTGCACTACCCCAAGAACTTCGACAACGCCCACTCTGGCCACAACAAGCACACCGACATCGGCTCTCTGACCCTGCTGTTCACCCCTCAGTGGGGCCTGCAGCTGCTGTCTCCCATTCAGAAGTCTAAGTCTTGGCTGTGGGTGCAGCCCCGACCCGGCCACGCCGTGATCAACGTGGGCGACTCTCTGCGATTCCTGTCTGGCAAGCGACTGAAGTCTTGTCTGCACCGAGTGTACCCCACCGGCGAGGTGTACCAAGAGGAGGACCGATACTCTATCGCCTACTTCCTGCGACCCGAGTCTGCCGCCAACTTCGAGGACGTGGACGGCAAGGTGGTGTCTGCCAAGCGATGGCACGACGAGAAGTACGTGACCTACACCGAGCCCCACGAGAAGCAAGACCTGTCTAACATCCTGACCGGCGGCATGGATCAGATCCTGGCCTAA  (621 bp) |
| Codon optimized L-threonine aldolase from *Pseudomonas sp.* coding DNA (*PsLtaE*) for *Y. lipolytica* based on UniProt ID: O50584 | ATGACCGATCAGTCTCAGCAGTTCGCCTCTGACAACTACTCTGGCATCTGTCCCGAGGCCTGGGCCGCCATGGAGAAGGCCAACCACGGCCACGAGCGAGCCTACGGCGACGATCAGTGGACCGCCCGAGCCGCCGACCACTTCCGAAAGCTGTTCGAGACCGACTGTGAGGTGTTCTTCGCCTTCAACGGCACCGCCGCCAACTCTCTGGCCCTGTCTTCTCTGTGTCAGTCTTACCACTCTGTGATCTGTTCTGAGACCGCCCACGTGGAGACCGACGAGTGTGGCGCCCCCGAGTTCTTCTCTAACGGCTCTAAGCTGCTGACCGCCCGATCTGAGGGCGGCAAGCTGACCCCCGCCTCTATCCGAGAGGTGGCCCTGAAGCGACAAGACATCCACTACCCCAAGCCCCGAGTGGTGACCATCACCCAAGCCACCGAGGTGGGCTCTGTGTACCGACCCGACGAGCTGAAGGCCATCTCTGCCACCTGTAAGGAGCTGGGCCTGAACCTGCACATGGACGGCGCTCGATTTTCTAACGCCTGTGCCTTCCTGGGCTGTACCCCCGCCGAGCTGACCTGGAAGGCCGGCATCGACGTGCTGTGTTTCGGCGGCACCAAGAACGGCATGGCCGTGGGCGAGGCCATCCTGTTCTTCAACCGAAAGCTGGCCGAGGACTTCGACTACCGATGTAAGCAAGCCGGACAGCTGGCCTCTAAGATGCGATTCCTGTCTGCCCCTTGGGTGGGCCTGCTGGAGGACGGCGCCTGGCTGCGACACGCCGCCCACGCCAACCACTGTGCTCAGCTGCTCTCTTCTCTCGTGGCCGACATCCCCGGCGTGGAGCTGATGTTCCCCGTGGAGGCCAACGGCGTGTTCCTGCAGATGTCTGAGCCCGCCCTGGAGGCCCTGCGAAACAAGGGCTGGCGATTCTACACCTTCATCGGCTCTGGCGGCGCCCGATTCATGTGCTCTTGGGACACCGAGGAGGCCCGAGTGCGAGAGCTGGCCGCCGACATCCGAGCCGTGATGTCTGCCTAA  (1041 bp) |
| Codon optimized malonyl-CoA reductase C-terminus from *Chloroflexus aurantiacus* coding DNA (*CaMCR-C*) for *Y. lipolytica* based on UniProt ID: A9WIU3 | ATGGCCGACCTGTCTGCCACCACCGGCGCCCGATCTGCCTCTGTGGGCTGGGCCGAGTCTCTGATCGGCCTGCACCTGGGCAAGGTGGCCCTGATCACCGGAGGCTCCGCCGGCATCGGAGGACAGATCGGCCGACTGCTGGCCCTGTCTGGCGCCCGAGTGATGCTGGCCGCCCGAGACCGACACAAGCTGGAGCAGATGCAAGCCATGATTCAGTCTGAGCTGGCCGAGGTGGGCTACACCGACGTGGAGGACCGAGTGCACATCGCCCCCGGATGTGACGTGTCTTCTGAGGCTCAGCTGGCCGACCTGGTGGAGCGAACCCTGTCTGCCTTCGGCACCGTGGACTACCTGATCAACAACGCCGGCATCGCCGGCGTGGAGGAGATGGTGATCGACATGCCTGTGGAGGGCTGGCGACACACCCTGTTCGCTAACCTGATTTCTAACTACTCTCTGATGCGAAAGCTGGCCCCCCTGATGAAGAAGCAAGGCTCTGGCTACATCCTGAACGTGTCTTCTTACTTCGGCGGCGAGAAGGACGCCGCCATCCCCTACCCCAACCGAGCCGACTACGCCGTGTCTAAGGCCGGACAGCGAGCCATGGCCGAGGTGTTCGCCCGATTCCTGGGCCCCGAAATTCAGATCAACGCCATTGCCCCTGGCCCTGTCGAAGGCGACCGACTGCGAGGAACCGGCGAACGACCCGGCCTGTTCGCCCGACGAGCTCGACTCATCCTGGAGAACAAGCGACTGAACGAGCTGCACGCCGCCCTGATCGCCGCTGCCCGAACCGACGAGCGATCTATGCACGAGCTGGTGGAGCTGCTCCTGCCCAATGACGTGGCCGCCCTGGAACAGAACCCTGCCGCCCCCACCGCCCTGCGAGAGCTGGCCCGACGATTCCGATCTGAGGGCGACCCCGCCGCCTCTTCCTCTTCTGCCCTGCTGAACCGATCTATCGCCGCCAAGCTGCTGGCCCGACTGCACAACGGCGGCTACGTGCTGCCCGCCGACATCTTCGCCAACCTGCCCAACCCCCCCGACCCCTTCTTCACCCGAGCTCAGATCGACCGAGAGGCCCGAAAGGTGCGAGACGGCATCATGGGCATGCTGTACCTGCAGCGAATGCCCACCGAGTTCGACGTGGCCATGGCCACCGTGTACTACCTGGCCGACCGAGTGGTGTCTGGCGAGACCTTCCACCCCTCTGGCGGCCTGCGATACGAGCGAACCCCTACCGGCGGCGAACTCTTCGGCCTGCCCTCTCCCGAGCGACTGGCCGAGCTGGTGGGCTCTACCGTGTACCTGATCGGCGAGCACCTGACCGAGCACCTGAACCTGCTGGCTCGAGCCTACCTGGAGCGATATGGCGCCCGACAAGTGGTGATGATCGTGGAGACCGAGACCGGCGCCGAGACCATGCGACGACTGCTGCACGACCACGTGGAGGCCGGCCGACTGATGACCATCGTGGCCGGCGATCAGATCGAGGCCGCCATTGACCAAGCCATCACCCGATACGGACGACCCGGCCCCGTGGTGTGTACCCCCTTCCGACCCCTCCCTACCGTGCCCCTGGTGGGCCGAAAGGACTCTGACTGGTCTACCGTGCTGTCTGAGGCCGAGTTCGCCGAGCTGTGTGAGCATCAGCTGACCCACCACTTCAGAGTGGCCCGATGGATCGCCCTGTCTGACGGCGCTCGACTGGCTCTCGTCACCCCCGAGACCACCGCCACCTCTACCACCGAGCAGTTCGCCCTGGCCAACTTCATCAAGACCACCCTGCACGCCTTCACCGCCACCATCGGCGTGGAGTCTGAGCGAACCGCTCAGCGAATCCTGATCAACCAAGTGGACCTGACCCGACGAGCCCGAGCCGAGGAGCCTCGAGACCCCCACGAGCGACAGCAAGAGCTGGAGCGATTTATTGAGGCTGTGCTGCTGGTGACCGCTCCCCTGCCTCCCGAGGCCGACACTCGATACGCCGGCCGAATCCACCGAGGCCGAGCCATCACCGTGTAA (2025 bp) |
| Codon optimized aspartate decarboxylase from *Shigella flexneri* coding DNA (*SfADC*) for *Y. lipolytica* based on UniProt ID: Q0T870 | ATGATCCGAACCATGCTGCAAGGCAAGCTGCACCGAGTGAAGGTGACCCACGCCGACCTGCACTACGAGGGCTCTTGTGCCATCGACCAAGACTTCCTGGACGCCGCCGGCATCCTGGAGAACGAGGCCATCGACATCTGGAACGTGACCAACGGCAAGCGATTCTCTACCTACGCCATCGCCGCCGAGCGAGGCTCTCGAATCATCTCTGTGAACGGCGCCGCTGCCCACTGTGCCTCTGTGGGCGACATCGTGATCATCGCCTCTTTCGTGACCATGCCCGACGAGGAGGCCCGAACCTGGCGACCCAACGTGGCCTACTTCGAGGGCGACAACGAGATGAAGCGAACCGCCAAGGCCATCCCCGTGCAAGTGGCCTAA  (381 bp) |
| Codon optimized monoamine oxidase from *Escherichia coli* coding DNA (*EcMAO*) for *Y. lipolytica* based on UniProt ID: P46883 | ATGGGCTCTCCCTCTCTGTACTCTGCCCGAAAGACTACTCTGGCCCTGGCCGTCGCCCTGTCTTTCGCCTGGCAAGCCCCCGTGTTCGCCCACGGCGGCGAGGCCCACATGGTGCCCATGGACAAGACCCTGAAGGAGTTCGGCGCCGACGTGCAGTGGGACGACTACGCTCAGCTGTTCACCCTGATCAAGGACGGCGCCTACGTGAAGGTGAAGCCCGGCGCTCAGACCGCCATCGTGAACGGACAGCCCCTGGCCCTGCAAGTGCCTGTGGTCATGAAGGACAACAAGGCCTGGGTGTCTGACACCTTCATCAACGACGTGTTTCAGTCTGGCCTGGATCAGACCTTCCAAGTGGAGAAGCGACCCCACCCCCTGAACGCCCTGACCGCCGACGAGATCAAGCAAGCCGTGGAGATCGTGAAGGCCTCTGCCGACTTCAAGCCCAACACTCGATTCACCGAGATCTCTCTGCTGCCCCCCGACAAGGAAGCCGTGTGGGCCTTCGCCCTGGAGAACAAGCCCGTGGATCAGCCCCGAAAGGCCGACGTGATCATGCTGGACGGCAAGCACATCATCGAGGCCGTCGTGGACCTCCAAAATAACAAGCTGCTGTCTTGGCAGCCCATCAAGGACGCCCACGGCATGGTGCTGCTGGACGACTTCGCCTCTGTGCAGAACATCATCAACAACTCTGAGGAGTTCGCCGCTGCCGTGAAGAAGCGAGGCATCACCGACGCCAAGAAGGTGATCACCACCCCCCTGACCGTGGGCTACTTCGACGGCAAGGACGGCCTGAAGCAAGACGCCCGACTGCTGAAGGTGATCTCTTACCTGGACGTCGGCGACGGCAACTACTGGGCCCACCCCATCGAGAACCTGGTGGCTGTGGTGGACCTGGAGCAAAAGAAGATCGTGAAGATCGAGGAGGGCCCCGTGGTGCCCGTGCCCATGACCGCCCGACCCTTCGACGGCCGAGATCGAGTGGCCCCCGCCGTGAAGCCCATGCAGATCATCGAGCCCGAGGGCAAGAATTACACCATCACCGGCGACATGATCCACTGGCGAAACTGGGACTTCCACCTGTCTATGAACTCTCGAGTGGGCCCCATGATCTCTACCGTCACCTACAACGACAACGGCACCAAGCGAAAGGTGATGTACGAGGGCTCTCTGGGCGGCATGATCGTGCCCTACGGCGACCCCGACATCGGCTGGTACTTCAAGGCCTACCTGGACTCTGGCGACTACGGCATGGGCACCCTGACCTCTCCCATCGCCCGAGGCAAGGACGCCCCCTCTAACGCCGTGCTGCTGAACGAGACCATCGCCGACTACACCGGCGTGCCCATGGAGATCCCCCGAGCCATCGCCGTGTTCGAGCGATACGCCGGCCCCGAGTACAAGCACCAAGAGATGGGACAGCCCAACGTGTCTACCGAGCGACGAGAGCTGGTGGTGCGATGGATCTCCACCGTGGGCAACTACGACTACATCTTCGACTGGATCTTCCACGAGAACGGCACCATCGGCATCGACGCCGGCGCCACCGGCATCGAAGCCGTGAAGGGCGTGAAGGCCAAGACCATGCACGACGAGACCGCCAAGGACGACACCCGATACGGCACCCTGATCGACCACAACATCGTGGGCACTACTCATCAGCACATCTACAACTTCCGACTGGACCTGGACGTGGACGGCGAGAACAACTCTCTGGTGGCCATGGACCCCGTGGTGAAGCCCAATACTGCCGGCGGCCCCCGAACCTCTACCATGCAAGTGAATCAGTACAACATCGGCAACGAGCAAGACGCCGCTCAGAAGTTCGACCCCGGCACCATCCGACTGCTGTCTAACCCCAACAAGGAGAACCGAATGGGCAACCCCGTGTCTTATCAGATCATCCCCTACGCCGGCGGCACCCACCCCGTGGCCAAGGGCGCTCAGTTCGCCCCCGACGAGTGGATCTACCACCGACTGTCTTTCATGGACAAGCAGCTGTGGGTGACCCGATACCACCCCGGCGAGCGATTCCCCGAGGGAAAGTACCCCAACCGATCTACCCACGACACCGGCCTGGGACAGTACTCTAAGGACAACGAGTCTCTGGACAACACCGATGCCGTCGTCTGGATGACCACCGGCACCACCCACGTGGCTCGAGCCGAGGAGTGGCCCATCATGCCCACCGAGTGGGTGCACACCCTGCTGAAGCCCTGGAACTTCTTCGACGAGACCCCCACCCTGGGCGCCCTGAAGAAGGACAAGTAA (2274 bp) |

**Table S3. Primer sequences used in this study.**

| Primer names | Sequences (5'--3') | Purposes |
| --- | --- | --- |
| pFBA-F | gcatccctaaatttgatgaaagcctaggCGTAATATTCCCCACCAAGCCGCTAGATA | pYL31 |
| pFBA-R | tcactcagatgcatagcacgcgtgtagaggatccGTGTAGTTTAGATTTCGAATCTGTGGG |  |
| Pex20t-F | cgtgctatgcatctgagtgaggtaccgactagtGGAAGTGTGGATGGGGAAGT |  |
| Pex20t-R | ctcctccgttattgtctcgctagctaactaGCTATATTTGACGATTGACG |  |
| pTDH1-F | gcatccctaaatttgatgaaagcctaggACGATCTACAGCCCGATCACATGAAC | pYL24 |
| pTDH1-R | tcactcagatgcatagcacgcgtgttctagaTGTTGATGTGTGTTTAATTC |  |
| Mig1t-F | cgtgctatgcatctgagtgaggtaccgactagtCACTGGCCGGTCGATAATTT |  |
| Mig1t-R | ctcctccgttattgtctcgctagctaactagatttAAACCCAAAAGGGCCGAAGG |  |
| Lip1-up-F | gcatccctaaatttgatgaaagcctaggTCTCTGATTCTCTTCTCCATTGTTATTTC | pUrLp-ΔLip1 |
| Lip1-up-R | cgtataatgtatgctatacgaagttatcctagTTAGTAGACAACAATCAGAACATCTCCC |  |
| Lip1-dn-F | cggccgcgcatgcaagtcgacCCAGTTTGAGATCCTAGACAAGTCTC |  |
| Lip1-dn-R | atgttacatccttttatcagacatagtcgcctaggATCATGAGACCCAACATTGTCATG |  |
| Dga1-up-F | gcatccctaaatttgatgaaagcctaggATGCTGCGGGCGGATCCTGGTGC | pUrLp-ΔDga1 |
| Dga1-up-R | cgtataatgtatgctatacgaagttatcctagAGCTTTTGTTTTGTGTGACTTG |  |
| Dga1-dn-F | cggccgcgcatgcaagtcgacGGAAAACTGCCTGGGTTAGGC |  |
| Dga1-dn-R | atgttacatccttttatcagacatagtcgcctaggTCTGATGGCCTGGAGCGAGTTTC |  |
| HBD1-up-F | gcatccctaaatttgatgaaagcctaggCTATCCTGAAATGTCCCATGTTCTC | pUrLp-ΔHBD1 |
| HBD1-up-R | cgtataatgtatgctatacgaagttatcctaCCACAGTTATAAGTATAAGATTG |  |
| HBD1-dn-F | cggccgcgcatgcaagtcgacTTGTGAAGCGAATGGGCAAGAT |  |
| HBD1-dn-R | atgttacatccttttatcagacatagtcgcctaggCTCTGTTTACTTGGTCTGCTCAGTACT |  |
| 2PS-F | accagcactttttgcagtactaaccgcagGGGTCATATAGCAGTGATGAT | pYLXP'-Gh2PS |
| 2PS-R | caggccatggaactagtcggtaccTTAGTTGCCGTTGGCCACTGC |  |
| ACC1-F | accagcactttttgcagtactaaccgcagCGACTGCAATTGAGGACACTA | pYLXP'-YlACC1 |
| ACC1-R | caggccatggaactagtcggtaccTCACAACCCCTTGAGCAGCTC |  |
| IboH-F | tcttgaattaaacacacatcaacaATGCCCCCCACCATCCGAAACAAC | pYL24-AmIboH |
| IboH-R | gatgcatagcacgcgtgtTCTAGATTAGGCCAGGATCTGATCCATGCCGC |  |
| LtaE-F | cagattcgaaatctaaactacacATGACCGATCAGTCTCAGCAG | pYL31-PsLtaE |
| LtaE-R | catagcacgcgtgtagaGGATCCTTAGGCAGACATCACGGCTC |  |
| MCR-C-F | accagcactttttgcagtactaaccgcagGCCGACCTGTCTGCCACCAC | pYLXP'-CaMCR-C |
| MCR-C-R | caggccatggaactagtcggtaccTTACACGGTGATGGCTCGGC |  |
| ADC-F | cagattcgaaatctaaactacacATGATCCGAACCATGCTGCAAG | pYL31-SfADC |
| ADC-R | catagcacgcgtgtagaggatccTTAGGCCACTTGCACGGGGAT |  |
| MAO-F | tcttgaattaaacacacatcaacaATGGGCTCTCCCTCTCTGTAC | pYL24-EcMAO |
| MAO-R | gatgcatagcacgcgtgttctagaTTACTTGTCCTTCTTCAGGGC |  |
| Ura-checkin-F | cagccaagaaaaccaacctgtgtgcttctctggatg | Check integration |
| Ura-checkin-R | catccagagaagcacacaggttggttttcttggctg |  |
| Lip1-checkout-F | atgaagactggcgtgtcgtcctttggtgacattggt | Check integration |
| Lip1-checkout-R | tcgccaatgatattggtatcaacgacggccccttca |  |
| Dga1-checkout-F | cagtggcactgatagaggaactaagaaacctc | Check integration |
| Dga1-checkout-R | gtctgagtctcttgacgttggaagaactcacc |  |
| Hbd1-checkout-F | agtgtaaaagcacggccattcagaatgtga | Check integration |
| Hbd1-checkout-R | taatagctctgttggcattttgctcactgc |  |

**Table S4** Strains were constructed in this research.

| Strain | Parental strain | Genotype or relevant properties | Source |
| --- | --- | --- | --- |
| Po1f (ATCC-MYA2613) | - | MatA, Leu2-270, Ura3-302, Xpr2-322, Axp1-2 | ATCC |
| Po1fk(--) | Po1f(--) | Δku70::loxP, (Leu^-^, Ura^-^) | [2] |
| TAL(-+) | Po1fk(--) | ΔLip1::P_TEFin_-*Gh2PS*-xpr2t, loxP-Ura3-loxP, (Leu^-^, Ura^+^) | This work |
| TAL(++) | TAL(-+) | ΔA08::Leu2, (Leu^+^, Ura^+^) | This work |
| TAL-ACC(-+) | TAL(--) | ΔDga1::P_TEFin_-*YlACC1*-lip2t, loxP-Ura3-loxP, (Leu^-^, Ura^+^) | This work |
| TAL-ACC(++) | TAL-ACC(-+) | ΔA08::Leu2, (Leu^+^, Ura^+^) | This work |
| TAL-ILM(-+) | TAL(--) | ΔDga1::P_TDH1_-*AmIboH*-mig1t, P_FBA_-*PsLtaE*-pex20t, P_TEFin_-*CaMCR-C*-xpr2t, loxP-Ura3-loxP, (Leu^-^, Ura^+^) | This work |
| TAL-ILM(++) | TAL-ILM(-+) | ΔA08::Leu2, (Leu^+^, Ura^+^) | This work |
| TAL-AMM(-+) | TAL(--) | ΔDga1::P_FBA_-*SfADC*-pex20t, P_TDH1_-*EcMAO*-mig1t, P_TEFin_-*CaMCR-C*-xpr2t, loxP-Ura3-loxP, (Leu^-^, Ura^+^) | This work |
| TAL-AMM(++) | TAL-AMM(-+) | ΔA08::Leu2, (Leu^+^, Ura^+^) | This work |
| TAL-A(-+) | TAL(--) | ΔDga1::P_FBA_-*SfADC*-pex20t, loxP-Ura3-loxP, (Leu^-^, Ura^+^) | This work |
| TAL-A(++) | TAL-A(-+) | ΔA08::Leu2, (Leu^+^, Ura^+^) | This work |
| Po1fk-AMM(-+) | Po1fk(--) | ΔDga1::P_FBA_-*SfADC*-pex20t, P_TDH1_-*EcMAO*-mig1t, P_TEFin_-*CaMCR-C*-xpr2t, loxP-Ura3-loxP, (Leu^-^, Ura^+^) | This work |
| Po1fk-AMM(++) | Po1fk-AMM(-+) | ΔA08::Leu2, (Leu^+^, Ura^+^) | This work |
| TAL-ΔHBD1(-+) | TAL(--) | ΔHBD1::loxP-Ura3-loxP, (Leu^-^, Ura^+^) | This work |
| TAL-ΔHBD1(--) | TAL-ΔHBD1(-+) | ΔHBD1::loxP, (Leu^-^, Ura^-^) | This work |
| TAL-ΔHBD1-ACC(-+) | TAL-ΔHBD1(--) | ΔDga1::P_TEFin_-*YlACC1*-lip2t, loxP-Ura3-loxP, (Leu^-^, Ura^+^) | This work |
| TAL-ΔHBD1-ACC(++) | TAL-ΔHBD1-ACC(-+) | ΔA08::Leu2, (Leu^+^, Ura^+^) | This work |
| TAL-ΔHBD1-ILM(-+) | TAL-ΔHBD1(--) | ΔDga1::P_TDH1_-*AmIboH*-mig1t, P_FBA_-*PsLtaE*-pex20t, P_TEFin_-*CaMCR-C*-xpr2t, loxP-Ura3-loxP, (Leu^-^, Ura^+^) | This work |
| TAL-ΔHBD1-ILM(++) | TAL-ΔHBD1-ILM(-+) | ΔA08::Leu2, (Leu^+^, Ura^+^) | This work |
| TAL-ΔHBD1-AMM(-+) | TAL-ΔHBD1(--) | ΔDga1::P_FBA_-*SfADC*-pex20t, P_TDH1_-*EcMAO*-mig1t, P_TEFin_-*CaMCR-C*-xpr2t, loxP-Ura3-loxP, (Leu^-^, Ura^+^) | This work |
| TAL-ΔHBD1-AMM(++) | TAL-ΔHBD1-AMM(-+) | ΔA08::Leu2, (Leu^+^, Ura^+^) | This work |

**Table S5. Comparison of the highest TRY metrics of engineered strains producing malonyl-CoA derivatives.**

| Product | Strain | Medium | Titer  (mg/L) | Yield  (mg/g DCW) | Productivity  (mg/L/h) | Related |
| --- | --- | --- | --- | --- | --- | --- |
| TAL | TAL | YPD | 1621.3 | 85.0 | 13.5 | Fig 2C |
| TAL | TAL-ACC | YPD | 2431.1 | 126.0 | 20.3 |  |
| TAL | TAL-ILM | YPD | 2638.8 | 138.1 | 22.0 |  |
| TAL | TAL-AMM | YPD | 2669.6 | 141.9 | 22.2 |  |
| 3-HP | TAL-ACC | YPD | 831.8 | 119.7 | 6.9 | Fig 4C |
| 3-HP | TAL-ACC | YPD+34mM β-Ala | 3042.0 | 35.4 | 25.4 |  |
| HHEP | TAL-AMM | YPD+34mM Asp | 5191.3 | 547.7 | 43.3 | Fig 4D |
| HHEP | TAL-AMM | YPD+34mM Asp +10Mm 3-HP | 6359.1 | 703.4 | 53.0 |  |
| TAL | TAL-A | YPD | 751.0 | 65.0 | 15.6 | Fig 4E |
| TAL | TAL-AMM | YPD | 407.9 | 54.9 | 8.5 |  |
| HHEP | TAL-A | YPD | 542.4 | 46.9 | 11.3 |  |
| HHEP | TAL-AMM | YPD | 3415.1 | 459.9 | 71.1 |  |
| TAL | TAL-ILM | YPD | 2211.4 | 106.0 | 18.4 | Fig 5A |
| TAL | TAL-ILM | YPD+34mM Glu | 2772.7 | 142.1 | 23.1 |  |
| HHEP | TAL-ILM | YPD | 99.9 | 6.2 | 2.1 |  |
| HHEP | TAL-ILM | YPD+34mM Glu | 113.9 | 6.8 | 1.6 |  |
| 3-HP | TAL-ILM | YPD | 1726.2 | 82.7 | 14.4 |  |
| 3-HP | TAL-ILM | YPD+34mM Glu | 2221.0 | 113.8 | 18.5 |  |
| TAL | TAL-AMM | YPD | 1182.8 | 79.4 | 9.9 | Fig 5C |
| TAL | TAL-AMM | YPD+34mM Asp | 1388.5 | 96.9 | 11.6 |  |
| HHEP | TAL-AMM | YPD | 3140.9 | 210.8 | 26.2 |  |
| HHEP | TAL-AMM | YPD+34mM Asp | 3601.1 | 250.9 | 30.0 |  |
| 3-HP | TAL-AMM | YPD | 1130.4 | 75.7 | 9.4 |  |
| 3-HP | TAL-AMM | YPD+34mM Asp | 1757.4 | 122.9 | 14.6 |  |
| HHEP | TAL-ΔHBD1-AMM | YPD | 128.4 | 6.5 | 1.8 | Fig 5E |
| HHEP | TAL-ΔHBD1-AMM | YPD+60g/L peptone | 314.8 | 18.2 | 6.6 |  |
| TAL | TAL-ΔHBD1-ACC | YPD | 2368.9 | 116.5 | 19.7 | Fig 5G |
| TAL | TAL-ΔHBD1-ILM | YPD | 2401.4 | 117.1 | 20.0 |  |
| TAL | TAL-ΔHBD1-AMM | YPD | 2310.4 | 113.6 | 19.3 |  |
| TAL | TAL-ΔHBD1-ILM | YPD+34mM Glu | 2822.6 | 136.2 | 23.5 |  |
| TAL | TAL-ΔHBD1-AMM | YPD+34mM Asp | 2893.4 | 139.8 | 24.1 |  |
| Lipid | TAL-ΔHBD1-ACC | YPD | 2121.9 | 100.6 | 17.7 |  |
| Lipid | TAL-ΔHBD1-ILM | YPD | 2187.1 | 102.8 | 18.2 |  |
| Lipid | TAL-ΔHBD1-AMM | YPD | 2152.0 | 100.7 | 17.9 |  |
| Lipid | TAL-ΔHBD1-ILM | YPD+34mM Glu | 2329.0 | 110.3 | 19.4 |  |
| Lipid | TAL-ΔHBD1-AMM | YPD+34mM Asp | 2356.9 | 110.5 | 19.6 |  |
| TAL | TAL-AMM | YPD | 1903.2 | 134.0 | 17.6 | Fig 6 |
| TAL | TAL-AMM | YPD+100mM Asp | 1586.5 | 97.2 | 14.7 |  |
| HHEP | TAL-AMM | YPD | 3597.3 | 251.6 | 33.3 |  |
| HHEP | TAL-AMM | YPD+100mM Asp | 5368.6 | 332.3 | 49.7 |  |
| 3-HP | TAL-AMM | YPD | 2499.0 | 173.3 | 23.1 |  |
| 3-HP | TAL-AMM | YPD+100mM Asp | 2380.8 | 150.0 | 22.0 |  |
| Lipid | TAL-AMM | YPD | 2311.6 | 161.3 | 21.4 |  |
| Lipid | TAL-AMM | YPD+100mM Asp | 2703.7 | 166.7 | 25.0 |  |
| OCFA | TAL-AMM | YPD | 148.9 | 10.4 | 1.4 |  |
| OCFA | TAL-AMM | YPD+100mM Asp | 182.5 | 11.3 | 1.7 |  |
